# Interface-guided phenotyping of coding variants in the transcription factor RUNX1 with SEUSS

**DOI:** 10.1101/2023.08.03.551876

**Authors:** Kivilcim Ozturk, Rebecca Panwala, Jeanna Sheen, Kyle Ford, Nathan Payne, Dong-Er Zhang, Stephan Hutter, Torsten Haferlach, Trey Ideker, Prashant Mali, Hannah Carter

## Abstract

Understanding the consequences of single amino acid substitutions in cancer driver genes remains an unmet need. Perturb-seq provides a tool to investigate the effects of individual mutations on cellular programs. Here we deploy SEUSS, a Perturb-seq like approach, to generate and assay mutations at physical interfaces of the RUNX1 Runt domain. We measured the impact of 115 mutations on RNA profiles in single myelogenous leukemia cells and used the profiles to categorize mutations into three functionally distinct groups: wild-type (WT)-like, loss-of-function (LOF)-like and hypomorphic. Notably, the largest concentration of functional mutations (non-WT-like) clustered at the DNA binding site and contained many of the more frequently observed mutations in human cancers. Hypomorphic variants shared characteristics with loss of function variants but had gene expression profiles indicative of response to neural growth factor and cytokine recruitment of neutrophils. Additionally, DNA accessibility changes upon perturbations were enriched for RUNX1 binding motifs, particularly near differentially expressed genes. Overall, our work demonstrates the potential of targeting protein interaction interfaces to better define the landscape of prospective phenotypes reachable by amino acid substitutions.

## INTRODUCTION

Cancer is a disease associated with progressive loss of cell identity and gain of signals promoting inappropriate survival and proliferation. It is now well established that somatic DNA mutations, particularly in oncogenes and tumor suppressors, can drive tumor development and progression [1–4]. It is also increasingly evident that different somatic mutations in the same gene can have different associations with prognosis and therapeutic response. For example, KRAS G13 but not G12 mutant colorectal cancers are sensitive to treatment with cetuximab [5]. In lung cancer, the G13 mutation is associated with shorter overall survival than the G12 mutation [6]. In breast cancer, TP53 mutations within DNA binding motifs are associated with worse prognosis than those outside, but within DNA binding motifs, mutations at codon 179 and the R248W substitution are associated with significantly poorer prognosis than other substitutions [7]. This highlights the need to develop strategies for studying perturbations at a finer scale than gene knock-out or knockdown.

High-throughput mutagenesis is emerging as a powerful tool to probe the varying consequences of different amino acid substitutions across the length of a protein or protein domain, however it is currently limited to specific functional readouts, such as target protein abundance [8] or functional assays [9–11]. Studying the effects of genetic perturbations on cellular programs and fitness has been challenging using traditional pooled screens. Approaches such as ScalablE fUnctional Screening by Sequencing (SEUSS) [12] and Perturb-seq [13] measure the transcriptional consequences of perturbations ranging from whole gene knockout to amino acid substitutions [14] in single cells, the latter making it possible to gain insight into the differences that distinguish mutations at the level of cellular programs relevant to cancer progression. SEUSS has been used to study the consequences of functional domain deletions and hotspot mutations to MYC in human pluripotent stem cells [12]. Perturb-seq applied to study driver mutations in KRAS and TP53 revealed that these mutations span a continuum of function that is not necessarily predicted by mutation frequency in cancer cohorts [14]. While providing greater function insight, these sequencing-based methods are not yet scalable to exhaustive mutagenesis, necessitating selection of target mutations.

Individual proteins often have multiple functions within cells, mediated through interaction with different binding partners. Somatic mutations in driver genes are overrepresented at interaction interfaces [15–19], suggesting that examining the consequences of perturbing distinct protein interfaces could provide a useful abstraction of the phenotypic space reachable by individual amino acid substitutions. To explore this hypothesis, we sought to design a library of mutations perturbing distinct physical interfaces of RUNX1, a transcriptional master regulator of hematopoiesis implicated in multiple cancer types, then use the resulting transcriptional landscape to examine cancer-associated mutations.

Runt-related transcription factor 1 (RUNX1) has been implicated in multiple tumor types such as acute myeloid leukemia and breast cancer [20,21]. The protein encoded by RUNX1 forms the heterodimeric complex core-binding factor (CBF) together with CBFB and is thought to be involved in the development of normal hematopoiesis [22,23]. CBF binds to the core element of many enhancers and promoters through the Runt domain of RUNX1 [22,24]. Specifically, RUNX members (RUNX1–3) modulate the transcription of their target genes by recognizing the core consensus binding sequence 5-TGTGGT-3, or very rarely, 5-TGCGGT-3, within their regulatory regions via their Runt domain [25]; while CBFB, a non-DNA-binding regulatory subunit in the CBF complex, binds to the Runt domain and leads to increased DNA-binding affinity [26]. A variety of transcriptional co-regulatory proteins bind RUNX1 via the Runt domain and modulate CBF activity [27].

As RUNX1 is a pioneer transcription factor and master regulator, we hypothesized that mutations affecting its interactions with transcriptional cofactors would manifest as changes to the expression of multiple RUNX1 target genes and thus potentially result in functional diversity. We applied SEUSS to study perturbations to protein interaction interfaces of the RUNX1 Runt domain, which harbors the majority of recurrent cancer mutations. Towards this, we planned for a library of 117 variants, including wild-type (WT) and loss-of-function (LOF) controls (**Table 1**). Thus, we prioritized variants with the potential to perturb distinct interactions, and therefore generate distinct transcriptional readouts implicating different aspects of the RUNX1 regulon. We analyzed these readouts to identify functionally distinct groups of RUNX1 mutations, characterize their effects on cellular programs, and study the implications for cancer mutations.

**Table 1.** Description of 117 library elements.

| Count | Class | Detail |
| --- | --- | --- |
| 83 | perturbation | perturbs at least one RUNX1 interaction |
| 1 | WT | contains WT RUNX1, no mutation |
| 17 | negative control | 10 silent or 7 neutral mutations |
| 1 | LOF | contains GFP, no RUNX1 |
| 10 | positive control | 5 truncating or 5 core mutations |
| 5 | combination | contains two or three mutations |

## RESULTS

### An interface-guided Perturb-seq assay for coding variant phenotyping of transcription factor RUNX1

While somatic mutations in RUNX1 span the entire gene, the most recurrent mutations cluster in the Runt domain that binds DNA and includes binding sites for multiple protein partners, including CBFB. We used protein structures (**Figure 1a**) and template-based docking [28] to identify 83 amino acid residues in the RUNX1 Runt domain involved in physical interactions with at least one of 33 protein partners with structural data (**Methods**, **Figure 1b, Supplementary Figure 1**). These were used to design an open reading frame (ORF) mutation library to assess the impact of perturbation of various RUNX1 interactions.

**Figure 1.**
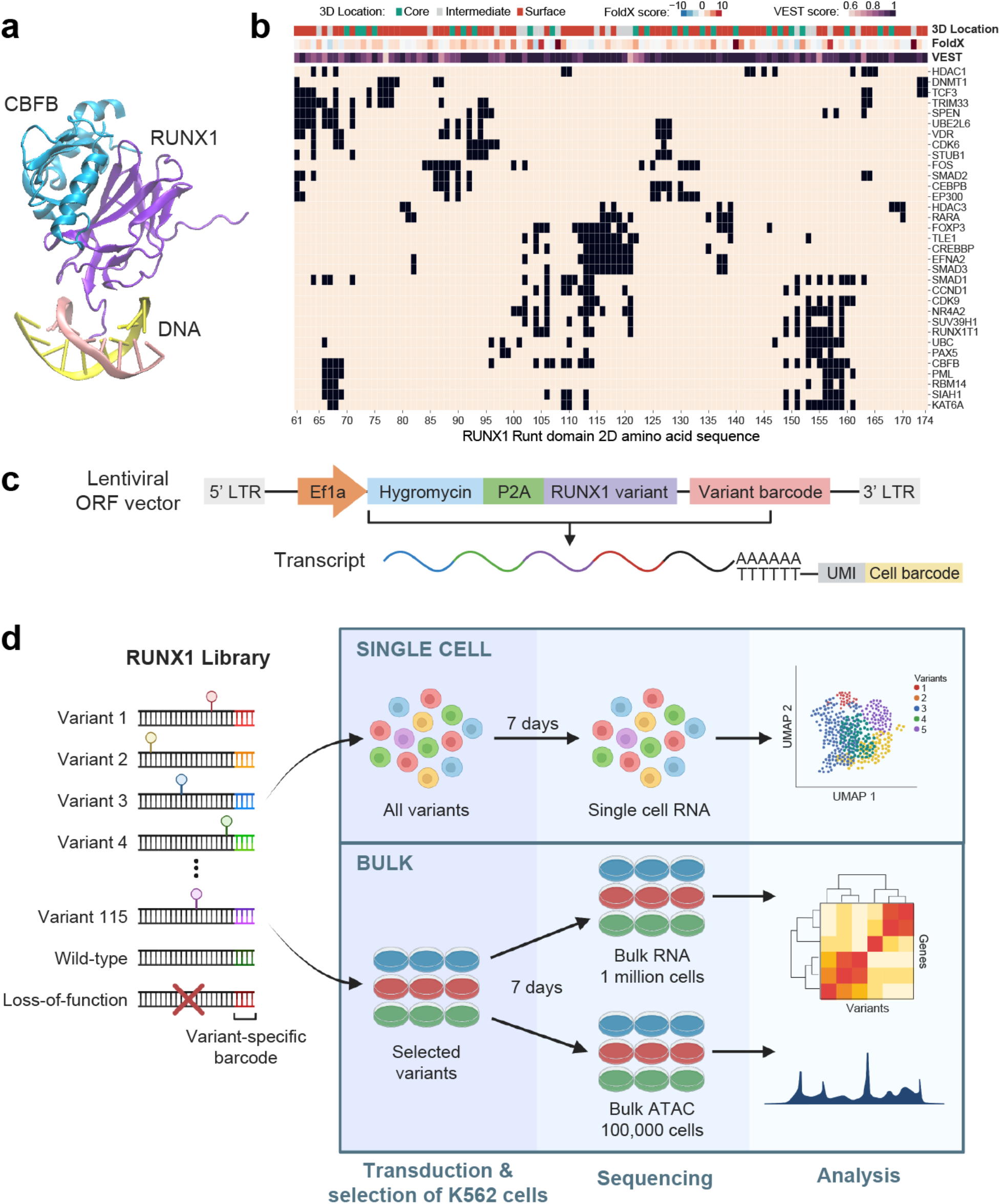
An interface-guided Perturb-seq assay for coding variant phenotyping of RUNX1. **a.** 3D crystal structure of transcription factor CBF, consisting of RUNX1 Runt domain (purple) and CBFB (blue), interacting with DNA (yellow and pink strands) (PDB: 1h9d). **b.** Amino acid residue map of the RUNX1 Runt domain. Columns represent amino acid residues, while rows represent interaction partners of RUNX1. At each row, interface residues involved in interaction to the partner are highlighted black. Rows are hierarchically clustered. On top: 3D location annotations of each residue (core, intermediate, and surface), followed by VEST and FoldX scores of most damaging mutations targeting the residue. The darker the color, the more damaging (VEST) or destabilizing (FoldX) the mutation is. **c.** Schematic of lentiviral ORF vector containing the RUNX1 variant (WT, mutated, or GFP) and a 12 base pair barcode sequence unique to each variant for identification during single cell transcriptome sequencing. **d.** Experimental and computational overview: ORF variant library design, transduction, single cell RNA-sequencing of all 117 library elements, bulk RNA and ATAC-sequencing of 12 selected library elements, and computational analysis.

For each of the 83 residues, we identified amino acid substitutions that would maximally perturb function based on pathogenicity scores from VEST [29] and folding free energy predictions from FoldX [30]. Where possible we also prioritized substitutions observed in tumors from the COSMIC database at these positions [31]. To provide a frame of reference for functional impact, we included WT RUNX1 and LOF controls (RUNX1 replaced with the green fluorescent protein (GFP)), along with 17 negative controls (expected to be indistinguishable from WT) consisting of 10 silent and 7 predicted neutral (based on VEST scores) mutations, and 10 positive controls (expected to have similar impact to LOF, by resulting in a truncated or unstable protein) consisting of 5 truncating and 5 core mutations. In order to evaluate the impact of multiple mutations, we also included 5 mutation combinations, bringing the total to 117 library elements (**Table 1**, **Supplementary Table 1**).

We used ScalablE fUnctional Screening by Sequencing (SEUSS) [12] to express the mutant RUNX1 library in K562 cells [32], with and without doxycycline-inducible CRISPRi knockdown of endogenous RUNX1 (**Methods**). Our library was generated from a modified lentiviral vector to contain a hygromycin resistance enzyme gene downstream of the EF1a promoter, followed by a P2A peptide motif, the RUNX1 variant (WT, mutated, or GFP as LOF control) and a 12 base pair barcode sequence unique to each variant for identification during single cell transcriptome sequencing (**Figure 1c**). The cells were transduced with the pooled variant library at a MOI of ∼0.3 to ensure that each cell received a single construct and were grown with hygromycin to select for cells carrying constructs. They were split into two populations, one treated with doxycycline to induce repression of the endogenous RUNX1 (dox), the other not (nodox). At day 7 post transduction, single cell RNA libraries were prepared and sequenced, with the remainder of cells being maintained until day 14 for fitness screening (**Figure 1d**).

We generated single-cell transcriptional profiles for 86,120 cells using the 10X Genomics Chromium platform [33], of which 48.4% contained detectable variant barcodes assigned to a single variant only. Four variants (G138V, S145I, P157R, T161I) were excluded due to low cell counts, and one negative control was removed (G143G) due to a frame shift artifact during the mutation library preparation. After additional quality control (QC) filtering, we recovered 40,522 high-quality cells with detectable variant barcodes assigned to a single variant only, covering 112 of 117 assayed variants for downstream analysis (**Methods**). 20,878 of these cells were from the pool treated with doxycycline (dox). These showed larger effect sizes associated with RUNX1 mutations relative to those still expressing endogenous RUNX1 (nodox) (**Supplementary Figure 2**). Therefore, we focused our analysis on the 20,878 high-quality cells without endogenous RUNX1 (median 136 cells per variant, **Supplementary Figure 3a, Supplementary Table 2**) (**Methods**). Among these, LOF (361 cells) and positive control variants (median 244 cells per variant) generated significantly higher numbers of cells in comparison to WT (127 cells) and negative control variants (median 63 cells per variant) (Pearson correlation; r=0.85, p<7.55e-09) (**Supplementary Figure 3b**), consistent with previous reports that reduction or loss of RUNX1 activates cell proliferation [34,35]. Mutation combinations had even higher numbers of cells than the LOF control (**Supplementary Figure 3b**).

We next analyzed variants according to their transcriptional profiles. We reasoned that variants with similar effects on RUNX1 targeting should cluster together, while those with distinct functional effects should cluster separately. To provide a frame of reference for functional impact, we compared expression profiles between cells harboring variants and cells carrying the WT or LOF control constructs. Variants were annotated as “WT-like” if the induced expression changes were indistinguishable from cells carrying the WT construct, and as “LOF-like” if indistinguishable from cells with the LOF construct. Any variant that did not have a WT-like expression profile was considered “functional”.

### Unsupervised clustering of cells and variants

After regressing out cell cycle effects, we performed unsupervised clustering of single cell gene expression profiles, which supported 3 clusters of cells (**Figure 2a**). Cluster 1 harbored the majority of cells expected to be functionally WT (the WT construct and negative controls; log(OR)=1.69, p<7.39e-19, and log(OR)=2.69, p<6.79e-301, respectively) (**Figure 2b**, **Supplementary Figure 4a**), whereas the LOF construct and positive control mutations were most enriched in clusters 2 and 3 (log(OR)=0.28, p<0.01 for LOF, and log(OR)=0.34, p<9.68e-15 for positive controls for cluster 2; and log(OR)=0.87, p<2.22e-15 for LOF, and log(OR)=0.88, p<2.54e-86 for positive controls for cluster 3) (**Figure 2b**, **Supplementary Figure 4a**). As expected, clusters did not show enrichment for cell cycle phases (**Supplementary Figure 4b-c**). Though cells containing perturbation variants were more enriched in cluster 1 overall (log(OR)=0.62, p<3.30e-82, **Supplementary Figure 4a**), certain variants were more concentrated in cluster 2 (e.g. S114L, log(OR)=1.28, p<4.76e-07, **Supplementary Table 3**), or cluster 3 (e.g. T169I, log(OR)=1.39 p<1.39e-15, **Supplementary Table 3**) (**Supplementary Figure 4e**).

**Figure 2.**
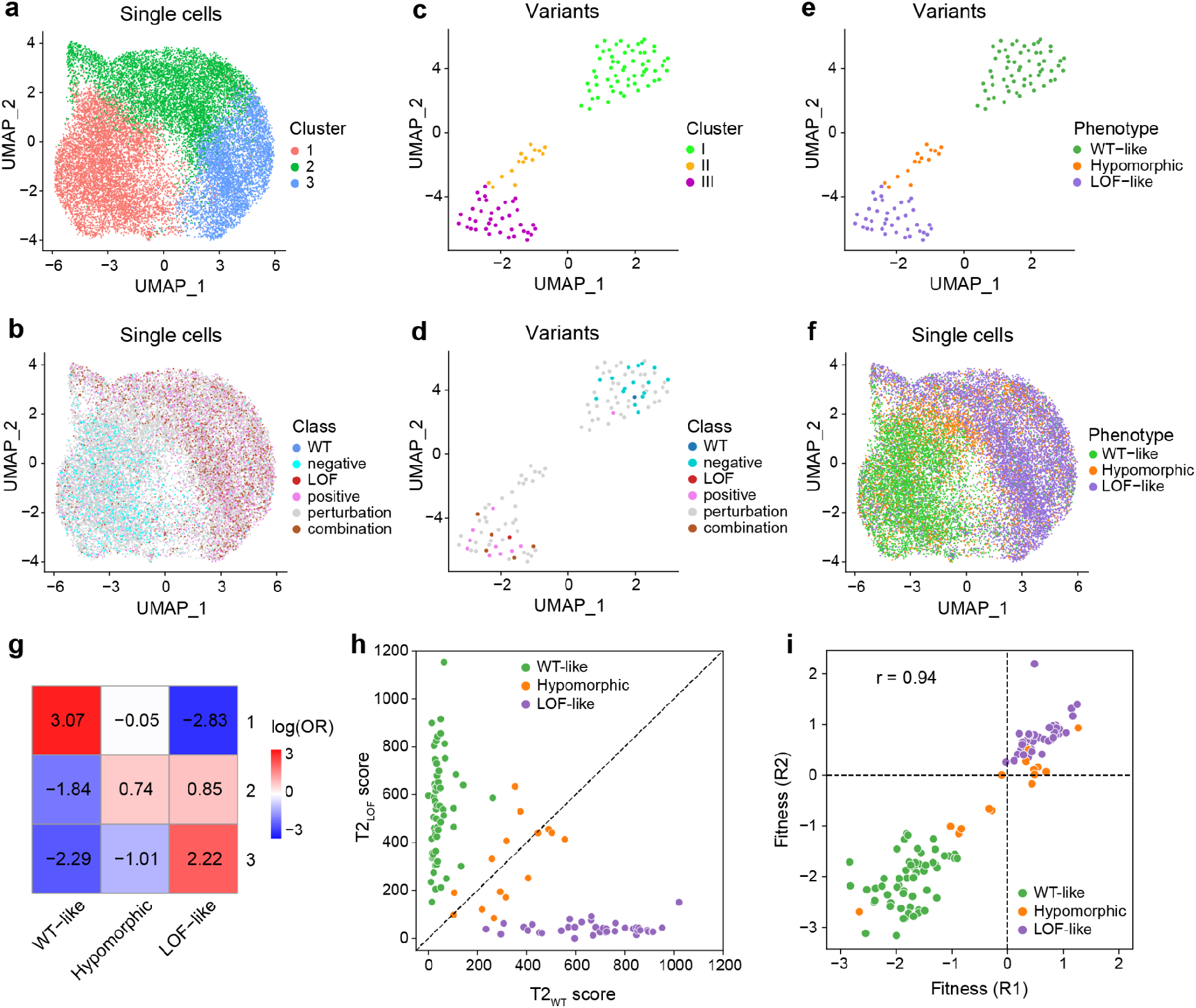
Unsupervised analysis of RUNX1 variant transcriptional effects informs WT-like, LOF-like and hypomorphic variants. **a-b.** UMAP embedding of single cells colored by **a.** unsupervised clusters, or **b.** variant classes, obtained using the top 2000 variable genes. Cell cycle effects are regressed out. **c-d-e.** UMAP embedding of variants constructed from mean gene expression vectors of the top 2000 variable genes for each variant, colored by **c.** unsupervised clusters, **d.** variant classes, or, **e.** variant functional designations of WT-like, LOF-like, and hypomorphic for unsupervised clusters in **c**. **f.** UMAP embedding of single cells colored by variant functional designations of WT-like, LOF-like, and hypomorphic obtained from unsupervised cluster designations in **e**. **g.** Cluster enrichment of single cells (unsupervised clusters from **a**) for assigned phenotypes (from **f**) based on log of odds ratios obtained using Fisher’s exact test. Positive values indicate enrichment, while negative values indicate depletion. **h.** T2 scores of each variant when compared to the WT (x-axis) or LOF (y-axis) control, colored by assigned phenotypes. **i.** Fitness scores of variants computed from two biological replicates, colored by assigned phenotypes (R1: replicate 1, R2: replicate 2).

To compare variants more directly, we averaged the expression of each of the top 2000 variable genes across all cells corresponding to each variant and performed unsupervised clustering based on these profiles, which again suggested three groups (**Figure 2c**). Group I included the WT construct and all negative control variants. Group III contained the LOF construct, and the majority of positive control variants (8 of 10) expected to result in a truncated or unstable protein (**Figure 2d**). Most of the perturbation variants also fell into these groups (41 variants into group I and 24 variants into group III), reflecting expression profiles similar to WT or LOF variants, whereas the separate assignment of 14 variants to group II suggested some partial loss of RUNX1 function, distinct from LOF or WT activity. Accordingly, we labeled variants in these groups as WT-like, hypomorphic, and LOF-like (**Figure 2e**).

In single cell space, cells carrying hypomorphic variants were most prevalent near the boundary between clusters 1 and 2 (**Figure 2f**) and were most enriched in cluster 2 (log(OR)=0.74, p<9.92e-89) (**Figure 2g**). Cell cluster 1 was highly enriched for cells harboring WT-like variants (log(OR)=3.07, p<2.22e-308), while cells harboring LOF-like variants were enriched both in clusters 2 (log(OR)=0.85, p<5.75e-172) and 3 (log(OR)=2.22, p<2.22e-308) (**Figure 2g**).

To further investigate how hypomorphic variants compared to the WT and LOF constructs, we quantified the differences in distributions of transcriptional profiles for groups of cells with the Hotelling’s two-sample T-squared test statistic (T2 score), a multivariate generalization of Student’s t-test [36]. Here we calculated T2 scores using the top 20 principal components across cells harboring each variant (**Methods**). Higher scores indicate a higher deviation from the reference variant (WT or LOF control). This statistical analysis revealed that all WT-like variants are indistinguishable from the WT construct via small T2 scores relative to WT (T2_WT_), but high T2 scores relative to the LOF control (T2_LOF_) (p<0.05, **Figure 2h**, **Supplementary Table 2**). Similarly, all LOF-like variants are indistinguishable from the LOF construct via small T2_LOF_ scores, but have high T2_WT_ scores (p<0.05, **Figure 2h, Supplementary Table 2**). Hypomorphic variants are significantly different from both the WT and LOF controls (p<0.05, **Figure 2h**, **Supplementary Table 2**) suggesting that these variants result in transcriptional changes that are not simply an intermediate between LOF and WT. This result is supported by differential gene expression analysis where 48 of the 141 genes differentially expressed between hypomorphic variants versus the WT control, were not differentially expressed between LOF versus WT, and 107 of the 232 genes differentially expressed between hypomorphic variants versus the LOF control, were also unique, suggesting gain of new activity (**Supplementary Figure 5**).

Our single cell data were derived from the pooling of two biological replicates, each of which was evaluated for cell fitness at day 14 (**Methods**). Fitness measurements were highly concordant between replicates (Pearson correlation; r=0.94, p<5.32e-55) (**Figure 2i**), were positively correlated with variant T2_WT_ scores (Pearson correlation; r=0.85, p<1.25e-32), and negatively correlated with T2_LOF_ scores (Pearson correlation; r=-0.77, p<8.40e-24). In general, LOF-like variants tended to be associated with increased fitness and larger cell numbers (**Supplementary Figure 3c**).

### Gene expression programs distinguishing RUNX1 variants

Hierarchical clustering of the top 2000 variable genes across 112 RUNX1 Runt domain variants (**Figure 3a**) once again separated WT-like, LOF-like and hypomorphic variants, and highlighted clusters of genes with similar expression patterns across each variant group. While dendrograms for all three groups showed additional substructure, we were particularly interested in the hypomorphic variants which appeared to separate into 3 sub-groups, which we referred to as hypomorphic-I, -II and -III, to reflect the progression of gene expression changes observed across the variants (Phenotype_5group, **Figure 3a**). Functional enrichment analysis of gene clusters (**Supplementary Figure 6, Supplementary Tables 4 and 5**) suggested genes involved in positive regulation of T cell lineage commitment, proliferation and activation were more highly expressed by WT-like variants (program 4), while genes associated with heme biosynthesis (program 1), angiogenesis (program 2) and regulation of the extracellular matrix (program 3) were more highly expressed in LOF-like variants. This is consistent with RUNX1’s role in hematopoietic lineage commitment and differentiation [22,23,37–39], with loss of RUNX1 leading to erythroid differentiation [40,41], and hematological malignancies [37,42]. The hypomorphic group showed higher expression of genes associated with tau protein kinase activity (program 9), which has been suggested to link nerve growth factor (NGF) to activation of mitogen-activated protein kinase (MAPK) signaling [43], but lowest expression of genes associated with neuronal plasticity and exocytosis (program 7), suggesting effects on some of RUNX1’s less-known functions [44–46].

**Figure 3.**
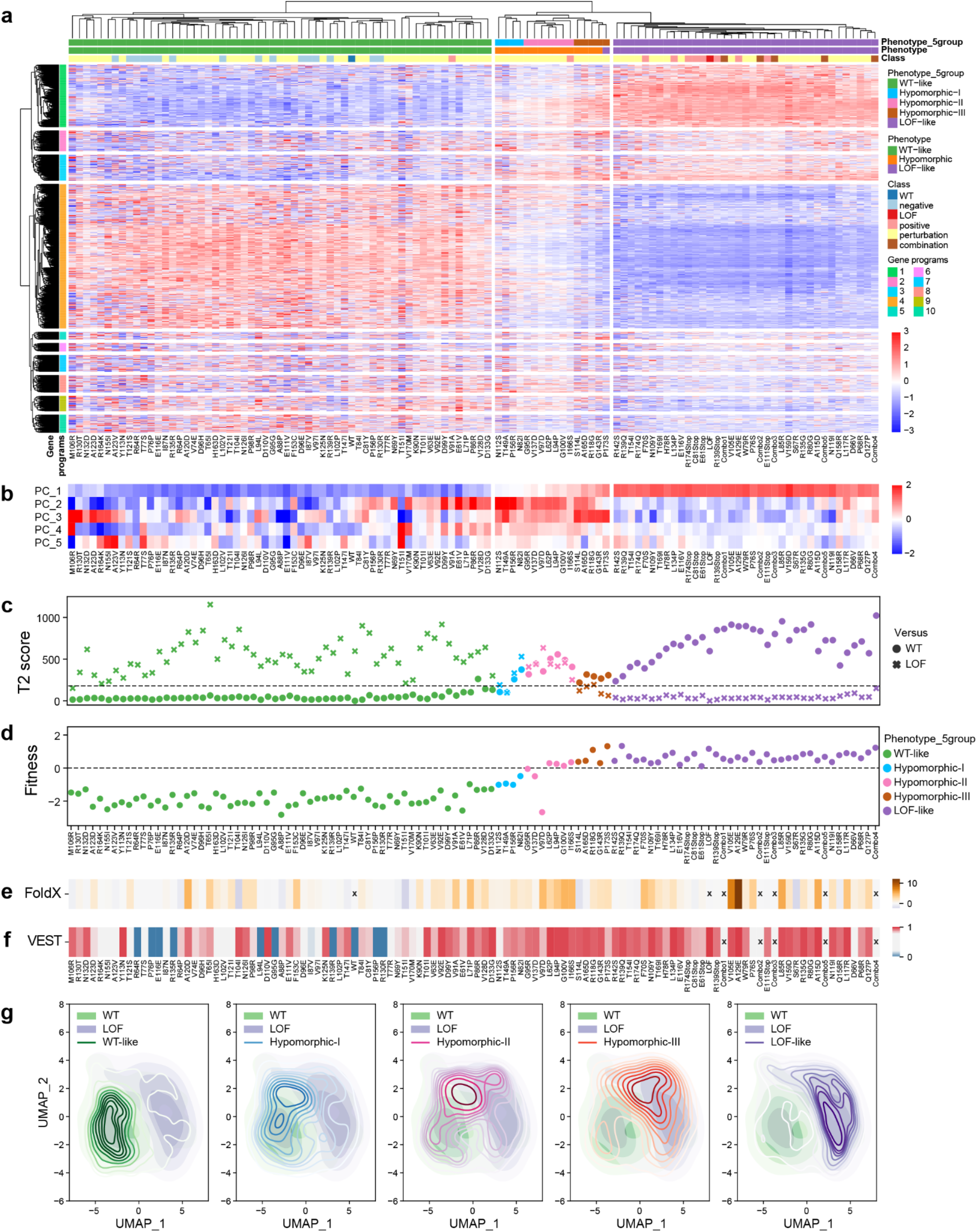
Mapping the phenotypic consequences of RUNX1 interface variants with transcriptomic analysis. **a.** Heatmap showing mean expression profiles of all variable genes (rows) in each variant (columns). Genes and variants are hierarchically clustered into ten (row colors) and five clusters (column colors: green: WT-like, blue: hypomorphic-I, pink: hypomorphic-II, brown: hypomorphic-III, and purple: LOF-like), respectively. The leaves of the variant dendrogram are ordered by increasing T2_WT_ scores. Gene expression values are z-scored. **b.** Top 5 PCs of each variant based on their mean gene expression profiles. Rows are scaled to have a mean of zero and unit variance. **c.** T2 scores of each variant when compared to the WT (circle) or LOF (cross) control, colored by phenotypes. Dotted line highlights value 178.79, the median of T2_WT_ scores of all WT-like and LOF-like variants. **d.** Mean fitness scores of variants, colored by phenotypes. **e.** FoldX, and **f.** VEST scores of each variant. Variants that could not be scored (combination of perturbation mutations, and/or WT and LOF variants) are grayed out and marked with an X. **g.** Kernel density estimate plots comparing UMAP embedding distributions of single cells belonging to each assigned phenotype (WT-like (green line), hypomorphic-I (blue line), hypomorphic-II (pink line), hypomorphic-III (brown line), and LOF-like (purple line)), to the cells overexpressing the WT (green shade) or LOF variant (purple shade).

The majority of variance in gene expression fell along the WT-like to LOF-like axis (PC1, 31.6% variance explained), both visually (**Figure 3b**), and statistically in the form of positive correlation with T2_WT_ scores (r=0.90, p<2.2e-16), and fitness scores (r=0.93, p<2.2e-16). On the other hand, PCs 2, 3 and 4 (3.4%, 3.1%, and 2.6% variance explained, respectively) pointed to transcriptomic effects more specific to the hypomorphic variants (**Figure 3b**). In particular, variants in the hypomorphic II group appear to have larger T2_WT_ and T2_LOF_ scores (**Figure 3c**) than the other hypomorphic variants, though this did not appear to affect fitness scores (**Figure 3d**).

Cell states can be described by the activities of sets of coordinated gene expression programs [47–51]. Analysis of single cell expression data with CoGAPS [52], a non-negative matrix factorization-based approach to infer groups of coregulated genes, identified 7 transcriptional patterns (**Supplementary Figure 7, Supplementary Table 6**). Patterns 4 and 1, most active in cells harboring WT-like variants, were enriched for the immune system, and cell cycle related functions respectively; while pattern 3, most active in cells harboring LOF-like variants, was enriched for heme biosynthesis pathways, consistent with a shift toward erythroid differentiation with loss of RUNX1 activity [40,41] . Cells harboring LOF-like variants were also more likely to show activity of patterns 6 and 7, which are enriched for stress response genes, and endoplasmic reticulum (ER) stress specifically, possibly suggesting that some variants may result in accumulation of misfolded RUNX1 protein in the ER. Patterns 2 and 5 were more active in cells harboring hypomorphic variants (**Supplementary Figure 7**). Pattern 2 interestingly is enriched for G1/S cell cycle and senescence, which could be indicative of G1 cell cycle arrest for some hypomorphic variants. Additionally, pattern 5, upregulated in various WT-like and hypomorphic variants, was enriched for interleukin signaling and cytokine signaling, suggesting a shift towards an inflammatory state for some cells [53–57].

LOF-like variants tended to have higher FoldX scores and VEST scores (**Figures 3e,f**). VEST scores were more strongly correlated with T2_WT_ scores and fitness scores than FoldX scores (T2_WT_: r=0.42, p<4.88e-06 vs r=0.38, p<6.31e-05; fitness r=0.43, p<3.23e-06 vs r=0.27, p<4.06e-03) suggesting that amino acid characteristics associated with pathogenicity are more likely to cause larger deviations from WT gene expression programs.

Kernel density estimates of single cell UMAP embeddings for variant groups demonstrate that the majority of cells belonging to each assigned phenotype (WT-like, hypomorphic-I, -II, -III, and LOF-like) occupy discrete regions in UMAP, though hypomorphic distributions seemed to harbor cells that also overlapped more closely with regions dominated by WT-like and LOF-like cells (**Figure 3g**). This could potentially be due to small differences in expression of the mutant construct, variability in knockdown of endogenous RUNX1 expression, stochasticity in measurement of gene expression, or even variable penetrance at the cellular level due to buffering built into cellular systems, such as stress response pathways.

### LOF-like variants significantly target RUNX1-DNA binding

Next, we looked at the distribution of RUNX1 perturbation variants along the 2D amino acid sequence of the Runt domain, annotated according to T2 score and functional effect (WT-like, LOF-like or hypomorphic) and noted some local clustering of variants with similar effect (**Figure 4a**). This clustering was also reflected in the 3D structural distribution of variants, especially in the region binding to DNA (**Figure 4b**). To further explore this, we investigated the impact of mutations at the interface of protein-nucleotide interactions of RUNX1. Eleven amino acid residues of RUNX1 (R80, R135, R139, R142, G143, K167, T169, V170, D171, R174, R177) were involved in

**Figure 4.**
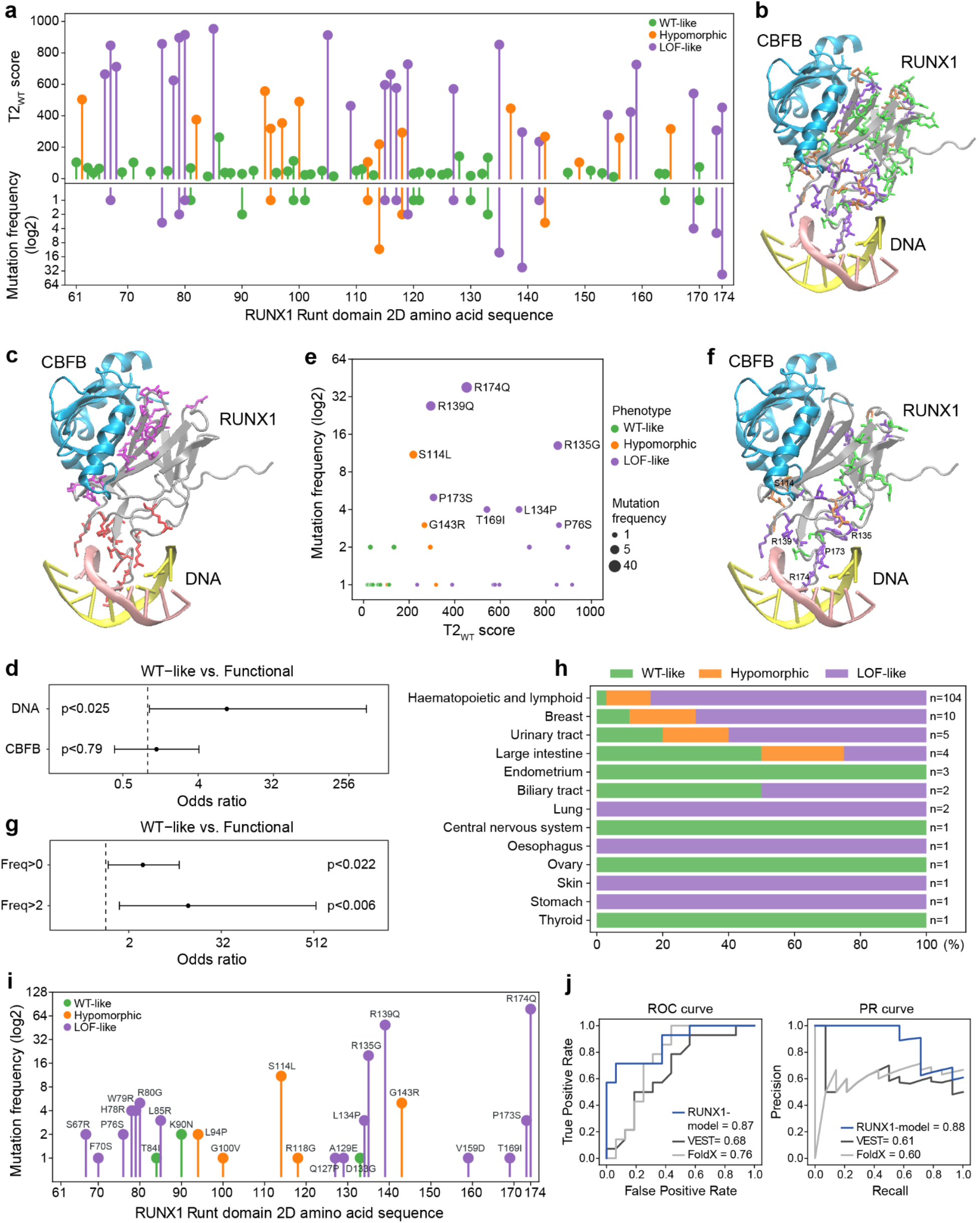
Mapping oncogenic variants into the RUNX1 regulatory landscape. **a**. Sequence-based phenotypic profiling of 79 RUNX1 perturbation variants. Variant T2 scores when compared to the WT (top panel), and mutation frequency in cancer (COSMIC) (log2 scaled) (bottom panel), distributed across 2D amino acid sequence of RUNX1 Runt domain, colored by phenotypes. **b-c-f.** Structure-based phenotypic profiling of RUNX1 perturbation variants. 3D crystal structure of transcription factor CBF, consisting of RUNX1 Runt domain (gray) and CBFB (blue), interacting with DNA (yellow and pink strands) is displayed (PDB: 1h9d). Amino acid residues corresponding to **b.** all 79 perturbation variants, **c.** only variants targeting DNA or CBFB interaction, or **f.** only variants observed in cancer are shown. **b-f.** Amino acid residues are colored based on phenotypic effects of corresponding variants (green: WT-like, orange: hypomorphic, purple: LOF-like), and 5 most frequent cancer mutations are annotated. **c.** Amino acid residues involved in interaction with DNA and CBFB are colored red and purple, respectively. **d.** Odds ratios (OR) and 95% confidence intervals using Fisher’s exact test. Enrichment or depletion of WT-like vs. functional (LOF-like or hypomorphic) impact variants for DNA or CBFB binding residues of RUNX1. OR>1 means enrichment for functional variants, while OR<1 means depletion. **e.** Scatter plot of library variants present in cancer in terms of T2_WT_ scores vs. mutation frequency (log2 scaled). Variants are colored based on their phenotypic annotations. **g.** Odds ratios and 95% confidence intervals using Fisher’s exact test. Enrichment or depletion of WT-like vs. functional impact variants for cancer mutations based on frequency. OR>1 means enrichment for functional variants, while OR<1 means depletion. **h.** A stacked bar plot showing percent distribution of phenotypic annotations of variants (WT-like, hypomorphic, and LOF-like) across cancers observed in different primary tissues. Sample size (n) for each tissue is displayed on the right. **i.** Frequency of missense mutations in the MLL dataset overlapping the variants in the RUNX1 library (log2 scaled). **j.** Performance of a classifier trained to predict WT-like and functional labels derived from transcriptional profiles (RUNX1-model), contrasted with VEST and FoldX scores. Performance is summarized by the area under the Receiver Operating Characteristic (auROC) and Precision-Recall (auPR) curves.

DNA binding based on the 3D crystal structure of the interaction (**Methods**), among which 8 residues are perturbed in our mutation library (R80G, R135G, R139Q, R142S, G143R, T169I, V170M, R174Q) (**Figure 4c**). These 8 mutations were significantly enriched for LOF-like impact vs. WT-like (Fisher’s exact test; OR=12.80, p<0.0084) and for functional (LOF-like or hypomorphic) vs. WT-like impact (Fisher’s exact test; OR=8.82, p<0.025, **Figure 4d**), consistent with the interruption of DNA-binding being very damaging to RUNX1-based gene regulation.

In comparison, for the 19 residues mediating the interaction with CBFB (**Methods**, **Figure 4c**), the enrichment for functional effects was much smaller; 10 of the mutations had LOF-like or hypomorphic impact while 9 had WT-like consequences (Fisher’s exact test; OR=1.27, p<0.79, **Figure 4d**). This may suggest that the DNA binding interface is more sensitive to amino acid sequence changes than protein interaction interfaces.

### Comparing coding variant Perturb-seq with recurrence in cancer

In principle, positions with functional variants that improve fitness would be selectively more advantageous for tumor cells and therefore more frequently mutated across patients. We tested whether variant-associated cellular phenotypes correlated with mutation frequency in cancer (COSMIC [31]). Frequency was weakly positively correlated with fitness (Pearson correlation; r=0.33, p<1.24e-03), but not with T2_WT_ scores (Pearson correlation, r=0.153, p<0.14, **Figure 4e**).

Nonetheless, functional mutations were significantly overrepresented among the subset of cancer mutations (Fisher’s exact test; LOF-like: OR=4.23, p<0.0084; functional (LOF-like or hypomorphic): OR=3.05, p<0.022, **Figure 4g**), consistent with positive selection for somatic mutations perturbing RUNX1 function in cancer. This impact was even more significant for cancer mutations with higher prevalence (frequency>2) (Fisher’s exact test; OR=14.87, p<0.0036; and OR=11.89, p<0.006, respectively, **Figure 4g**). Among the 5 most frequently observed mutations (**Figure 4e**), four of them (R174Q, R139Q, R135G and P173S) target amino acid residues involved in DNA binding and display LOF-like impact; while the remaining one (S114L) targets a residue involved in CBFB binding and is hypomorphic (**Figure 4f)**.

Of the RUNX1 mutations shared between our assay and the COSMIC database (**Figure 4e**), the majority of occurrences were in hematopoietic malignancies (n=104), followed by breast cancer (n=10), the urinary tract (n=5) and the large intestine (n=4). Across these four tumor types, approximately 79.6% of observed variants were LOF-like and 14.6% were hypomorphic (**Figure 4h**). We revisited RUNX1 mutations in a larger set of hematopoietic malignancies from the Munich Leukemia Laboratory **(Methods)**. This dataset included 717 tumors with somatic missense mutations in the Runt domain, of which 201 tumors contained 24 unique variants present in our library **(Figure 4i)**. Once again, we observed a bias for LOF-like or any functional mutations (Fisher’s exact test; OR=14.20, p<4.36e-05; and OR=11.03, p<7.66e-05, respectively). Notably, the same mutations were most frequent in these tumors (R174Q, R139Q, R135G and S114L).

### Using transcriptome-based labels for variant functional classification

COSMIC and MLL datasets encompassed missense mutations that were not in our library, both substitutions at different positions and different amino acid substitutions at some of the included positions. We reasoned that our library could serve as training data for predicting functional effects of other Runt domain variants. The 79 perturbation variants (41 WT-like, 24 LOF-like and 14 hypomorphic) in our library were divided into a training and test set at random while preserving the ratio of functional and WT-like variants (**Methods**). We annotated each variant with 85 features from the SNVBox database [58] describing substitution effects on amino acid biophysical properties, evolutionary conservation of variant sites, local sequence biases, and site-specific functional annotations and assigned a class label of WT-like or functional based on transcriptional profiles. We then trained a Random Forest classifier using the training set, and tested on the test set, obtaining area under the curve scores of 0.87 for ROC (auROC) and 0.88 for Precision-Recall curves (auPR) (**Figure 4j**). This RUNX1-specific model outperformed sequence-based variant effect prediction from VEST and protein stability predictions from FoldX.

Encouraged by these results, we trained a classifier using all 79 perturbation variants and evaluated its performance using the positive and negative missense control variants included in our library, obtaining an auROC of 0.81 and an auPR of 0.84. We then predicted transcriptomic effect labels (functional or WT-like) for all remaining possible missense mutations on the RUNX1 protein (n=2582), resulting in an additional predicted 302 functional and 355 WT-like mutations in the RUNX1 Runt domain not contained in our library. For the subset of mutations observed in cancer (COSMIC or MLL) (**Supplementary Figure 8**), predictions were biased toward being functional (COSMIC: 101 functional vs. 52 WT-like, Fisher’s exact test: OR=2.93, p-value<1.96e-08; MLL: 109 functional vs. 36 WT-like, OR=5.00, p-value<1.01e-15). Predictions for all Runt domain variants are described in **Supplementary Table 7**.

### Hypomorphic variant impact on the RUNX1 regulon

To validate the hypomorphic effect variants, we performed bulk RNA and ATAC sequencing, and investigated whether they altered DNA accessibility at regulatory elements bound by RUNX1 near differentially expressed genes. We selected 12 variants to study, including the WT and LOF control variants and 9 hypomorphic variants that showed the largest deviation from control conditions: two hypomorphic-I (N82I, P156R), six hypomorphic-II (L62P, G95R, V97D, G100V, V137D, I166S) and one hypomorphic-III (R118G). We also included one LOF-like variant (V159D) predicted to be involved in RUNX1-CBFB binding to further investigate the effects of interrupting this interaction (**Table 2**). Three of the hypomorphic variants (G95R, G100V and R118G) were observed in human tumors (COSMIC or MLL) (**Figure 5a**, **Table 2**).

**Table 2.**
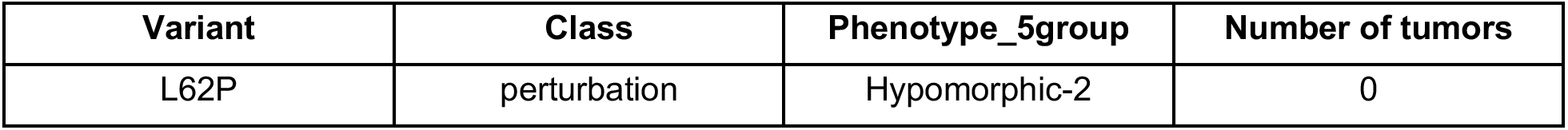
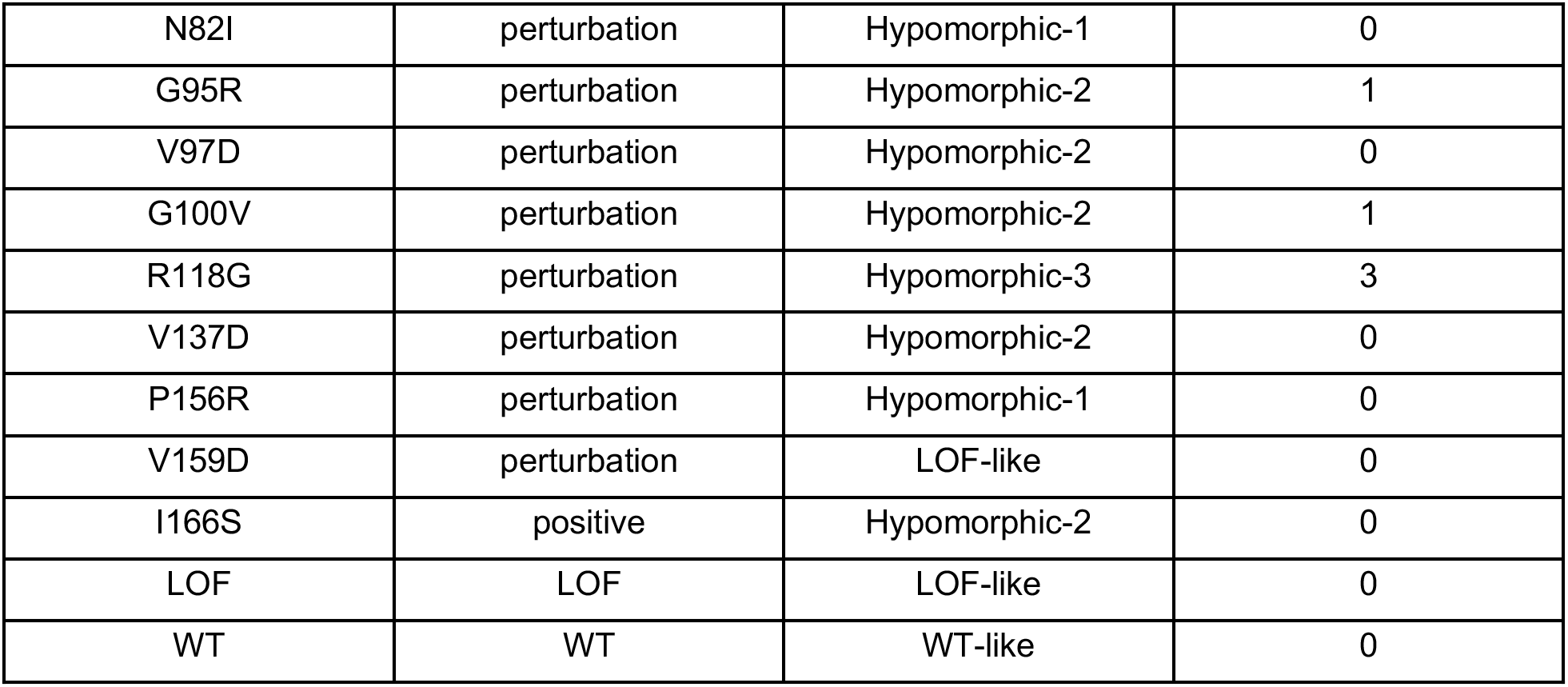
Twelve validation variants selected for bulk RNA- and ATAC-sequencing. Three variants were observed in human tumors in COSMIC or MLL cohorts.

**Figure 5.**
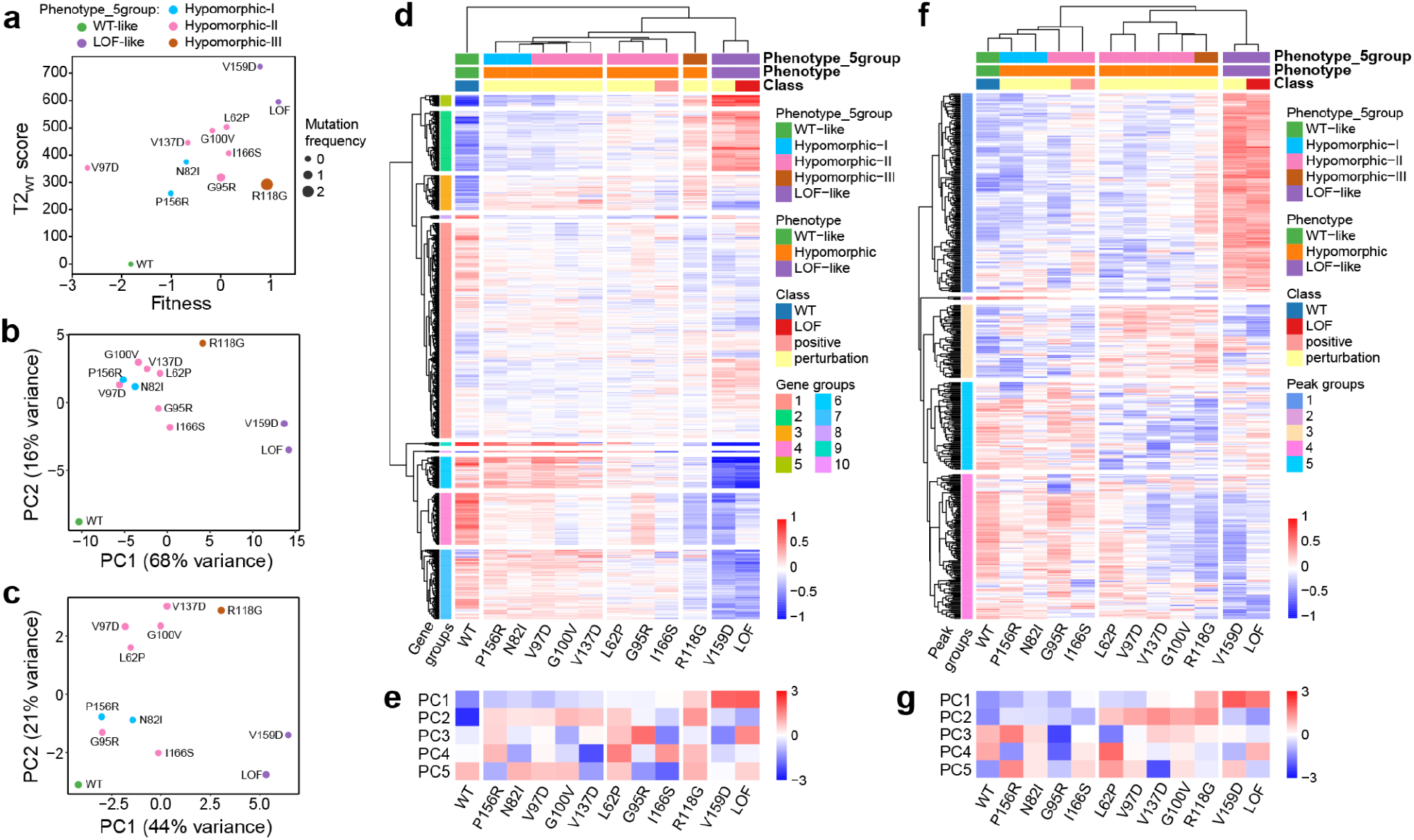
Bulk RNA- and ATAC-seq analysis of 12 validation variants. **a.** Overview of 12 validation mutations in terms of T2_WT_ and fitness scores (obtained from the scRNAseq analysis), and mutation frequency in cancer. Cancer mutations are annotated. **b.** PCA plot of 12 validation mutations, in the bulk RNA-seq setting, using top 2000 variable genes from scRNA-seq. Gene expression is averaged across replicates. **c.** PCA plot of 12 validation mutations, in the bulk ATAC-seq setting, using top 500 variable peaks. DNA accessibility is averaged across replicates. **d.** Hierarchical clustering of samples (rows) and genes (columns) in the bulk RNA-seq setting, using top 2000 genes obtained from scRNA-seq. Gene expression is averaged across replicates. The leaves of the variant dendrogram are ordered by increasing T2_WT_ scores. Gene expression values are mean-centered. **e.** Top 5 PC embeddings of each sample based on mean gene expression across replicates. Rows are z-scored. **f.** Hierarchical clustering of samples (rows) and peaks (columns) in the bulk ATAC-seq setting, using top 500 variable peaks. DNA accessibility is averaged across replicates. The leaves of the variant dendrogram are ordered by increasing T2_WT_ scores. DNA accessibility values are mean-centered. **g.** Top 5 PC embeddings of each sample based on mean DNA accessibility across replicates. Rows are z-scored.

We performed bulk RNA- and ATAC-seq screens for each variant separately in K562 cells grown in doxycycline to induce repression of the endogenous RUNX1, and hygromycin to select for transduced cells. The screen was conducted in three biological replicates per mutation with more than 1 million cells in each replicate. At day 7 post transduction, the cells were split into two groups: ∼1 million cells to be sequenced to a depth of 30 million reads/sample for bulk RNA-seq, and 100,000 cells to be sequenced to a depth of 75 million reads/sample for bulk ATAC-seq (**Figure 1d**).

After alignment, QC filtering, normalization, removal of replicate-specific batch effects, and averaging across replicates, we obtained an 18,646 gene by sample expression matrix, and an 82,870 peak by sample count matrix (**Methods**). Using the top 2000 variable genes from the scRNAseq analysis, we performed PCA, and hierarchical clustering of variants based on bulk gene expression. PC1 once again correlated with the progression of phenotypic effects (WT-like, hypomorphic-I, -II, -III, and LOF-like) (**Figure 5b**). PCA with the top 500 variable ATAC-seq peaks showed a similar trend (**Figure 5c**). Unsupervised hierarchical clustering on bulk gene expression (**Figure 5d**) reproduced the earlier scRNA-seq based clustering (**Figure 2a**), with the difference between WT and LOF conditions driving the main axis of variance (PC1, **Figure 5e**) and the second (PC2) distinguishing hypomorphic variants from both WT and LOF, with more specific distinctions captured by PCs 3 and 4. The similarity between bulk and single cell analysis supports that single cell transcriptomic analysis can reliably identify hypomorphic variants. Unsupervised hierarchical clustering of the top 500 ATAC-seq peaks produced a similar ordering of variants (**Figure 5f,g**), suggesting this peak set contained information relevant to variant-specific gene expression effects. We saw similar results when replicates were analyzed separately (**Supplementary Figure 9**).

To investigate whether hypomorphic variants altered DNA accessibility at RUNX1-bound regulatory elements near differentially expressed genes, we studied 202 genes that were significantly upregulated and 89 genes that were significantly downregulated in at least one hypomorphic variant relative to both WT and LOF controls (FDR<0.05, **Methods**). Of these, 63 and 27 respectively had ATAC peaks in their promoter regions ((-1 kb, +100 bp) of transcription start sites) that also contained RUNX1 binding motifs, suggesting direct regulation by RUNX1 (**Methods**). Hierarchical clustering based on the 63 upregulated genes clearly showed higher expression in hypomorphic variants (**Figure 6a**) while the same analysis for the 27 confirmed downregulation (**Figure 6b**). Analyzing these genes for functional enrichment (**Supplementary Table 8**) suggested hypomorphic variants may upregulate interleukin-10 signaling and PERK regulated gene expression (**Figure 6c**) but downregulate FGFR1 and IGFR1-regulated signaling (**Figure 6d**). To further evaluate specific genes, we visualized RNA and ATAC profiles for cells carrying hypomorphic mutations relative to cells with WT and LOF controls. As an example of a gene on a continuum from WT to LOF, PTPN22 shows an intermediate effect of the hypomorphic variant G100V, a variant we observed in the MLL dataset, on gene expression and DNA accessibility (**Figure 6e**). In contrast, CXCL2, involved in interleukin-10 signaling, and FGFR1, the main driver of fibroblast growth factor receptor 1 (FGFR1) signaling, demonstrated potential gain- or loss-of-function activity relative to both WT and LOF controls (**Figure 6f-g**). RUNX1 mutations are reportedly more frequent in the context of FGFR1 translocations which have been linked to more aggressive disease [59]. For FGFR1, we observed multiple ATAC peaks in the hypomorphic case that were not present in either the WT or LOF cases, possibly suggesting effects of the variant on targeting RUNX1 to motifs with inhibitory activity on gene expression.

**Figure 6.**
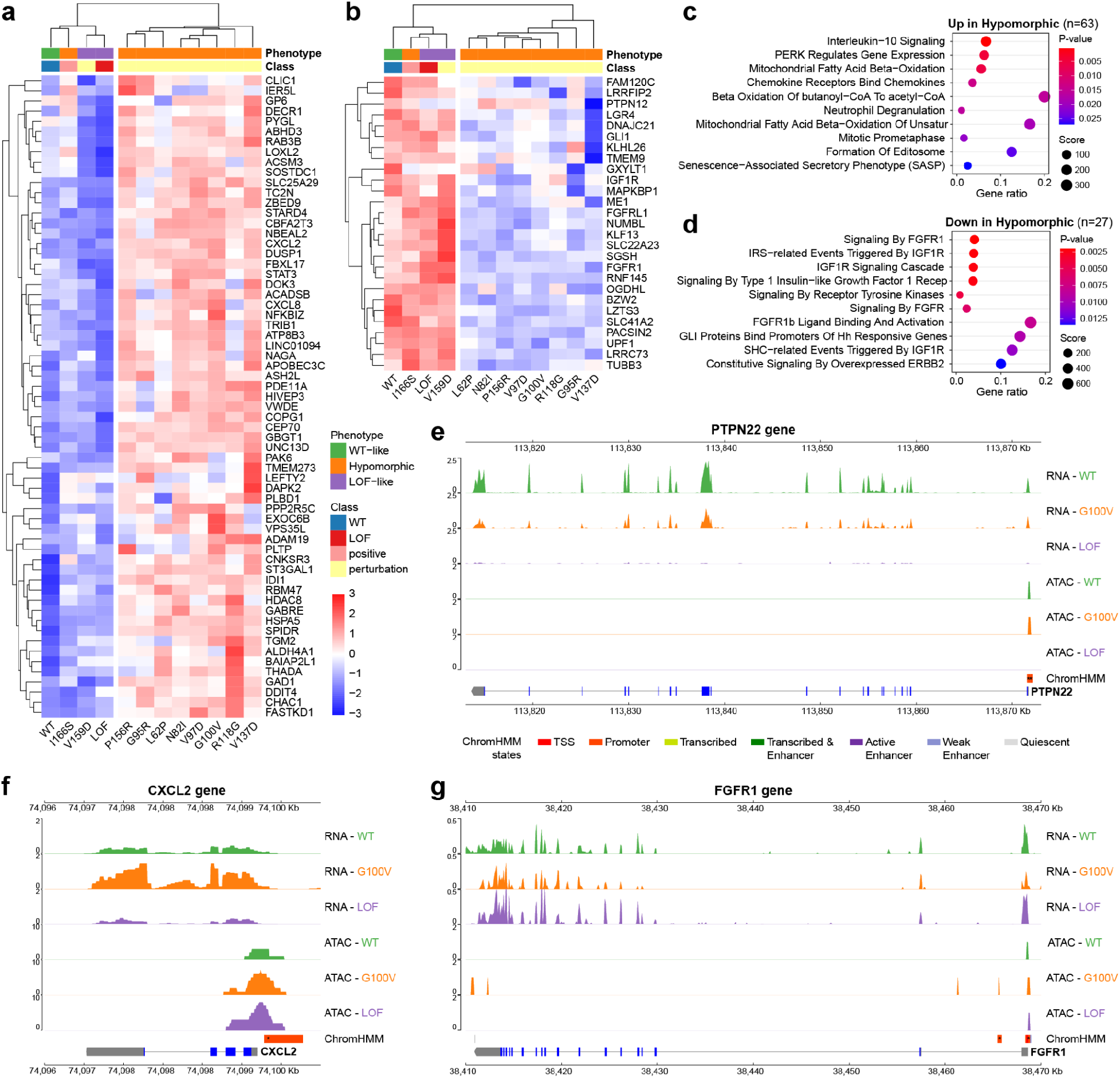
Regulatory consequences of hypomorphic RUNX1 Runt domain variants. **a-b.** Hierarchical clustering of samples (rows) and genes (columns) in the bulk RNA-seq setting, for genes that are directly regulated by RUNX1 and are **a.** upregulated (n=63), or **b.** downregulated (n=27), in at least one hypomorphic variant against both WT and LOF controls. Gene expression is averaged across replicates. Gene expression values are z-scored. **c-d.** Gene set overrepresentation analysis results for Reactome pathways for genes in panels **a** and **b**, respectively. Top 10 terms ordered by p-values are displayed. **e-f-g.** RNA-seq and ATAC-seq tracks of 3 example genes: **e.** NCF2 showing partial loss of function in hypomorphic against WT control, **f.** CXCL2 showing gain-of-function against both WT and LOF controls, and **g.** FGFR1 showing loss-of-function against both WT and LOF controls. Tracks are displayed for WT and LOF controls along with the G100V hypomorphic variant. ATAC-seq peaks are annotated with ChromHMM states, and star (*) indicates the presence of a RUNX motif. Exons and UTRs of each gene are displayed with blue and gray bands, accordingly.

Several additional differentially up- and down-regulated genes specific to cells with hypomorphic variants had RUNX1 motifs in nearby enhancers (**Supplementary Figure 10a-b**). Of these, several genes including STAT3 and MAPKBP1 had RUNX1-bound ATAC peaks both at its promoter and a nearby enhancer making it difficult to discern which element contributed to its altered expression. Upregulated genes showed enrichment for interleukin-10 signaling, similar to the promoter analysis (**Supplementary Table 9**). Additionally, they showed enrichment in NGF-stimulated transcription and NTRK1 regulated signaling, while downregulated genes were enriched in NFAT activation and BCR signaling (**Supplementary Figure 10c-d**). Available chromatin conformation capture data for K562 cells supported chromatin looping between RUNX1 bound enhancers and up and downregulated genes. For example, we observed loops linking enhancers to increased expression of CD24 for RUNX1 variant V137D, and to decreased expression of RPP25 for R118G, a variant previously observed in human tumors, relative to both WT and LOF controls (**Supplementary Figure 10e-f**). In both cases, ATAC profiling of the hypomorphic variant suggested the involvement of RUNX1 bound enhancers whereas ATAC peaks were not observed for WT and LOF controls, illustrating that hypomorphic variants can perturb both transcription enhancing and inhibitory functions of RUNX1 on gene regulation.

## DISCUSSION

While there is increasing appreciation that different mutations affecting the same cancer genes can lead to differences in disease severity [6,7,60] or drug sensitivity [5,61,62], the potential for pleiotropy to drive heterogeneous tumor cell phenotypes remains poorly understood. In this work we used information about physical contacts between proteins to guide design of a library of amino acid substitutions and profiled their transcriptional consequences at the single cell level using SEUSS. We selected RUNX1 as our target protein for its well-studied role in cancer and its activity as a master regulator of hematopoiesis, reasoning that point mutations in this gene could result in large detectable differences in gene expression. We focused our analysis on the Runt domain which harbors the majority of recurrent cancer mutations affecting RUNX1 and designed mutations based on *in silico* prediction of their potential to disrupt physical interfaces with transcriptional co-factors. Analysis of the resulting single cell transcriptomic profiles revealed that the majority of mutations we selected had effects similar to loss of function or wild type conditions excepting 15 variants that generated transcriptional profiles that differed from both extremes.

This study included two important updates to our SEUSS vector designs to improve signal in our screens: 1) to eliminate issues with barcode shuffling we positioned the variant and variant barcode in direct proximity; 2) to ensure minimal modification to the ORF we positioned the variants downstream of the 2A peptide thereby only a proline is appended to the N-terminus of the protein sequence. Simultaneous knockdown of endogenous RUNX1 further boosted effect sizes. Notably, the differences in transcriptional profiles of hypomorphic variants at the single cell level reproduced robustly in bulk RNA data, supporting SEUSS as a viable strategy for investigating the relatively subtle differences in gene expression we observed for missense variants.

The role of RUNX1 as an oncogene versus tumor suppressor is still not entirely clear and may depend on the type of malignancy as well as the other mutations present. In humans and in *vivo* model systems, loss of RUNX1 leads to increased susceptibility to AML; point mutations affecting RUNX1 are associated with shorter time to progression from MDS to AML and worse prognosis in AML and CML [63,64]. In contrast, in disease with RUNX1 translocations, a survival dependency on WT RUNX1 has been reported [65]. On the K562 background, loss of function appears to be associated with higher fitness, suggesting a more tumor suppressive role in this setting (CML with blast crisis), though it is important to note that K562 cells represent disease that developed on a WT RUNX1 background.

In this single cell experiment, the majority of mutations resulted in WT-like effects, with loss of function (LOF) variants being the second most common class. LOF mutations, and especially variants affecting DNA binding were associated with higher fitness scores, and larger cell counts suggesting that RUNX1 acts as a tumor suppressor in K562 cells, and its loss provides a selective growth advantage. Notably, the 5 constructs carrying pairs of mutations had an even more extreme effect than the LOF control that replaced RUNX1 with GFP.

A subset of hypomorphic variants showed subtle differences from the loss of function variants, which was confirmed by bulk RNA-seq data. Though small, these differences showed the potential to affect key signaling pathways (NGF-stimulated transcription, interleukin-10 signaling, PERK activity) and cancer genes (CBFA2T3, ETV4, FGFR1, GLI1, SGK1 and STAT3). Cells harboring hypomorphic variants were less commonly found to express a transcriptional program associated with neuronal plasticity and exocytosis while overexpressing genes associated with response to NGF. An increasing body of literature implicates neurotrophic signaling as a promoter of malignant cell growth and survival across a variety of tumor types [66]. As hypomorphic variants had lower expression of FGFR1 perhaps this indicates a different pattern of reliance on growth factors with links to neuronal plasticity [67]. Certain immune functions were also altered, for example IL-10 signaling molecules, CXCL2 and CXCL8, implicated in neutrophil recruitment [68], were higher in the hypomorphic case, whereas NFATC1, a mediator of T cell activity, was downregulated. The physiological implications of these small differences in the tumor microenvironment is unclear, though increased neutrophil-to-lymphocyte ratio has been associated with poor prognosis [69].

A number of the mutations in our library were recurrently observed across multiple tumors, a phenomenon usually associated with oncogenes [1], however the most recurrent events still clustered with loss of function controls. Several hypomorphic variants were also seen in multiple tumors, though they did not reach the same level of recurrence. In single cell plots, individual mutations were difficult to distinguish without first mapping to densities. Even then, it was apparent that individual cells could coincide with WT or LOF mutation regions. Further investigation is needed to understand whether this reflects stochastic differences in construct expression or knockdown of endogenous RUNX1, cell-to-cell differences in read coverage, or *bonafide* variable penetrance of the variant effect on the phenotype of individual cells.

Limitations of our study include that it was performed exclusively in K562 cells. It is unclear how well the effects will generalize to other leukemic cell lines or non-hematopoietic tumor types. In addition, we focused only on the Runt domain, whereas other domains are also important for RUNX1 interaction with cofactors. Thus, we may not have fully captured the space of possible phenotypes that can be generated by single amino acid substitutions in RUNX1. Furthermore, our epigenetic profiling was limited to DNA accessibility, whereas ChIP-seq would more directly reveal mutation-associated changes to RUNX1 localization. These questions will be the topics of future studies to better understand the role perturb-seq can play in providing exploitable mechanistic insights in cancer.

## CONCLUSION

Understanding the consequences of single amino acid substitutions in driver genes remains an unmet need. New technologies are making it possible to place such variants into the context of cellular programs and fitness. Our study demonstrates the potential of targeting protein interactions to explore the space of cellular phenotypes reachable by single amino acid substitutions.

## Acknowledgements

This work was supported by NIH grant 1U01HG012059 to HC, NIH grants R01HG012351, OT2OD032742, U54CA274502 and a Department of Defense Grant W81XWH-22-1-0401 to PM, NIH grant R01 DK098808 to DZ and CCMI pilot project funds under NIH Grant U54CA209891 to TI. Computational infrastructure support was provided by NIH grant P41GM103504. We thank Rafael Bejar, Adam Klie, Jennifer Phuong Nguyen, Agnieszka D’Antonio-Chronowska and Kelly Frazer for many helpful discussions.

## Author contributions

Original concept and project supervision by HC. Project planning, design, and method development by KO, PM, DZ and HC. Experiments by RP, JS, KF, NP and PM. MLL dataset preparation by SH and TH. Data acquisition, processing and analysis by KO and JS. Preparation of manuscript by KO, RP, JS, TI, PM, and HC.

## METHODS

### RUNX1 reference

Variant residue positions were defined based on the RUNX1B isoform of the RUNX1 gene (ENSG00000159216), corresponding to the Q01196-1 isoform protein (453 amino acids) described as the canonical sequence in the UniProt database [70], encoded by transcript ENST00000344691 (7274 base pairs) [71]. The Runt domain is ∼128 amino acids long, corresponding to amino acid positions 50-177 in the RUNX1 protein [72,73].

### Protein 3D structure analysis

We obtained 61 experimentally verified undirected protein interactions of RUNX1 with a confidence score higher than 0.4 from STRING v9.1 [74]. Experimental 3D co-crystal protein structures for RUNX1-CBFB interaction (PDB: 1ljm, 1e50, 1h9d) were obtained from the Protein Data Bank (PDB) [75], and used to predict amino acid residues of RUNX1 in direct physical contact with CBFB as described in our previous work [76]. The remaining interactions did not have co-crystal structures. Instead, we used *in silico* template-based protein docking on single protein structures with PRISM [28] to identify contact residues. PRISM returned predictions for 33 RUNX1 interaction partners (**Figure 1b**).

Amino acid residues of RUNX1 involved in DNA binding (PDB: 1h9d) were determined using the distance between two non-hydrogen atoms of amino acids and nucleotides, one from the protein and one from the DNA. If the distance was less than 3.5A, we designated those residues as interface residues [17]. Amino acid residues were annotated as core, surface, or intermediate based on their relative solvent accessible surface areas as described in our previous work [76]. VMD [77] was used to visualize protein 3D structures (**Figures 1a** and **4a,d**).

### Selection of variants for library construction

The ORF mutation library consists of 117 elements: 83 single amino acid substitutions at protein interaction interfaces in the RUNX1 Runt domain, 1 WT construct, 1 LOF construct, 17 negative, and 10 positive control mutations, and 5 combinations of two or more interface mutations (**Figure 1d**, **Table 1, Supplementary Table 1**). Variant effect prediction scores for all possible missense mutations targeting each residue were obtained from VEST [29] and FoldX [30]. Variant frequency in human tumors was determined from the COSMIC database [31] (obtained on 11/7/2022, for transcript ENST00000344691). For each residue, the most damaging amino acid substitution possible from a single base substitution (the highest VEST or FoldX scored mutation) was chosen to be included in the ORF mutation library, prioritizing cancer mutations where possible, to maximize the possibility of perturbing physical protein interactions. 30 of 83 mutations tested are cancer mutations.

The WT construct consists of WT RUNX1, while the LOF construct contains a green fluorescent protein (GFP) in place of RUNX1. 17 negative control mutations consist of 10 silent and 7 neutral (predicted based on VEST scores) mutations and are expected to be functionally indistinguishable from the WT construct. 10 positive controls consist of 5 truncating and 5 core mutations and are expected to have similar impact to the LOF construct. 5 perturbation mutation combinations consist of two or three combinations of perturbation mutations already in the library (**Table 1**, **Supplementary Table 1**).

### Experimental Methods

#### Building of RUNX1 variant library

The gene overexpression vector was generated from a modified lentiviral vector (Addgene #120426). The vector was modified by removing both the mCherry transgene and the hygromycin resistance enzyme gene. The hygromycin resistance enzyme gene was then re-cloned to be immediately downstream of the EF1a promoter, followed by a P2A peptide motif and a NheI restriction site, which was used to clone in the library elements. A 12 base pair barcode sequence was then introduced downstream of the cloning site to identify variants during single cell transcriptome sequencing. To insert the barcode, the backbone was digested with NheI (New England BioLabs), and a pool of 12 base pair long barcodes with flanking sequences compatible with the NheI site was cloned using Gibson assembly. To clone the library elements, the expression vector was digested with NheI for 3 hours at 37 °C. The linearized vector was then purified using a QIAquick PCR Purification Kit (Qiagen).

DNA fragments coding for the library elements were ordered from Twist Bioscience as a site saturation variant library in an arrayed format as linear dsDNA. A fraction of each oligonucleotide encoding the corresponding variant was then combined, and the pool was amplified via PCR using KAPA-Hifi (Kapa Biosystems) in 50 μL reactions containing 10 ng of pooled template and 2.5 μL of primers RX1_01 and RX1_02 (10 mM), which include ∼30 bp of DNA homologous to the overexpression vector to enable Gibson assembly cloning. A thermal cycler was used to heat the sample to 95 °C for 3 minutes, then 16 cycles of 98 °C for 20 seconds, 68 °C for 15 seconds, and 72 °C for 45 seconds, followed by a final 5 minute extension at 72 °C. The PCR products were then purified using Agencourt AMPure XP Beads (New England BioLabs) beads at a 0.8:1 bead:PCR reaction ratio. See Supplementary Table 10 for primer sequences.

Gibson assembly was then used to clone the pooled library elements into the overexpression vector. For the reaction, 50 ng of the digested vector and 30 ng of the insert were mixed with 5 mL of Gibson Reaction Master Mix (New England BioLabs) in a reaction volume of 10 μL. The Gibson reactions were incubated at 50 °C for 1 hour and transformed via heat shock into 50 mL of One Shot Stbl3 chemically competent cells (Invitrogen). This was done by incubating the cells with the Gibson on ice for 30 minutes, followed by a 45 second heat shock at 42 °C then 2 minutes on ice, then the addition of 250 μL of SOC media (Thermo Fisher Scientific). The cells were allowed to recover shaking at 37 °C for 1 hour and were then plated on LB-carbenicillin plates. Individual bacterial colonies were picked off of the plate and grown in LB-carbenicillin culture media shaking for 16 hours at 37 °C. After growth, plasmid DNA was isolated via a Qiagen Plasmid Mini Kit. Each colony was Sanger sequenced using the primer RX1_03 to identify the variant, then by the primer RX1_04 to capture the associated barcode. One overexpression vector was created for each variant, each with a single unique barcode associated. After ∼30% of the library was cloned, the oligonucleotides for remaining elements were re-pooled and cloned using the above protocol, until the full library was assembled. To generate the combination mutations, the first mutation was created as described above. Subsequent mutations were generated with overlap extension PCR with primers containing the desired mutations.

#### Lentivirus production

Replication deficient lentiviral particles were produced in an arrayed format for each element of the library in HEK293FT cells (Invitrogen) via transient transfection. The HEK293FT cells were grown in DMEM media (Gibco) supplemented with 10% FBS (Gibco) and 1% antibiotic-antimycotic (Thermo Fisher Scientific). One day prior to transfection, HEK293FT were plated in 12 well plates at ∼35% confluency, giving one well per element of the library. The day of transfection, the culture medium was removed and replaced with fresh DMEM plus 10% FBS. Meanwhile, the transfection mix was prepared by mixing 125 μL of Optimem reduced serum media (Life Technologies) with 1.5 μL of lipofectamine 2000 (Life Technologies), 125 ng of pMD2.G plasmid (Addgene #12259), 500 ng of pCMV delta R8.2 plasmid (Addgene #12263), and 375 ng of each plasmid overexpression vector for each library element. The transfection mix was incubated for 30 minutes, then added dropwise to the HEK293FT cells. The viral particles in the supernatant were harvested at 48 and 72 hours post transfection, and the virus for each library element were pooled and filtered with a 0.45 mm filter (Steriflip, Millipore), then concentrated to 1.5 mL using Amicon Ultra-15 centrifugal filters with a cutoff 100,000 NMWL (Millipore). The virus was then mixed, aliquoted and frozen at -80 °C. For the validation screen, the transfection was performed in 15 cm dishes, one for each of the selected validation mutations, and frozen separately.

#### Generation of clonal inducible RUNX1 repression cell line

To repress the endogenous RUNX1, the repression vector was generated from a PiggyBac inducible dCas9 construct (Addgene #63800). The vector was modified by removing the inducible transgene, the sequence for the KRAB-dCas9 fusion (Addgene #60954) followed by a P2A sequence then GFP was inserted in its place. The vector was then modified through the insertion of a U6 promoter followed by SaII and AflII cloning sites for insertion of guide RNA sequences, then a guide RNA scaffold. Guides for CRISPRi targeting RUNX1 were chosen from the Dolcetto library set A [78] and ordered via oligonucleotide from IDT. The guides were then cloned into the repression vector after digestion with SaII (New England BioLabs) and AflII (New England BioLabs).

K562 cells (ATCC) were cultured in RPMI 1640 media (Gibco) supplemented with 10% FBS and 1% antibiotic-antimycotic. One day prior to electroporation, the K562 cells were maintained at a concentration of 1 million cells per mL. The day of the electroporation, the cells were spun down and resuspended at a concentration of 10 million cells per mL. A total of 2 μg of DNA was added to 100 μL of cells containing a 1:2.5 molar ratio of the all-in-one RUNX1 targeting repression vector to the PiggyBac transposase vector (Transposagen). The DNA was then electroporated into the K562 cells using the Ingenio Electroporation Kit (Mirus Bio) and a 4D Nucleofector (Lonza) per the manufacturer’s protocol. The cells were recovered for 3 days, then selected for those that received integration by the addition of 1 μg/mL puromycin (Gibco) into the culture media. After 4 days of selection, the cells were split across a 96-well plate into single colonies by serial dilution. Individual colonies were then grown and assessed for their degree of inducible RUNX1 repression.

#### Quantification of RUNX1 expression

To measure RUNX1 repression in the single colonies, each colony was split into two separate populations and grown in RPMI media supplemented with 10% FBS and 1% antibiotic-antimycotic and 1 μg/mL puromycin. In one of the groups, 1 mg/mL doxycycline (Thermo Fisher Scientific) was added to the media to induce expression of the dCas9-KRAB transgene. Both sets of cells were maintained at 200,000 cells/mL over the course of 3 days after the addition of the doxycycline. On day 3 the cells were pelleted, and RNA was extracted using a Qiagen RNeasy Mini Kit. Complementary DNA (cDNA) was synthesized from the RNA using the Protoscript II First Strand cDNA Synthesis Kit (New England BioLabs) per the manufacturer’s protocol, then diluted 1:4 with water. To quantify expression, qPCR was performed on the cDNA using a CFX Connect

Real Time PCR Detection System (Bio-Rad). For each sample, two sets of primers were used; a set used to quantify RUNX1 expression (**Supplementary Table 10**) which was compared to the housekeeping gene GAPDH. The qPCR was carried out in a total volume of 10 μL containing 5 μL of iTaq Universal Sybr Green Master Mix (Bio-Rad), 2 μL of each primer (10 μM), and 1 mL of diluted cDNA. Thermal cycling conditions were 95 °C for 2.5 minutes, followed by 40 cycles of 95°C for 10 seconds, then 60 °C for 30 seconds. All samples were run in triplicate, and the RUNX1 expression was determined using the 2-delta delta CT method, by comparing to the GAPDH expression. The clone with the highest degree of RUNX1 repression was selected for use in the screen and subsequent experiments.

#### Sequencing screening

The K562 clonal cell line previously generated for repression of the endogenous RUNX1 protein was cultured in RPMI media supplemented with 10% FBS and 1% antibiotic-antimycotic. For the single cell RNA sequencing screen, the cells were transduced with the pooled variant library at an MOI of ∼0.3 to ensure that each cell received a single construct. The viral transduction was performed by mixing the virus with media containing 8 μg/mL polybrene (Millipore). The cells were suspended in this media at a concentration of 2 million cells per mL and spun at 1000 G for 2 hours at 33 °C in a 12-well plate. The cells were then pelleted and resuspended in fresh media at a concentration of 400,000 cells/mL. 24 hours after transduction, the media was again changed, and the cells were resuspended at 400,000 cells/mL. 48 hours after transduction, the cell culture media was changed to media containing 1 μg/mL puromycin and 200 μg/mL hygromycin (Invitrogen) to select for transduced cells. At that time the cells were also split into two separate populations, and to one of the populations doxycycline was added daily at a concentration of 1 μg/mL to induce repression of the endogenous RUNX1. Throughout the duration of the screen, the media was changed each day, and the cells were maintained at a concentration of 400,000 cells/mL. The screening was conducted with two biological replicates with greater than 1 million cells in each condition to ensure greater than 1000-fold coverage of the library. At day 7 post transduction, a subset of the cells was processed with single cell RNA sequencing, with the remainder of cells being maintained until day 14 for fitness screening.

For the bulk RNA sequencing and bulk ATAC sequencing screen, the cells were transduced with the twelve validation mutations separately. 48 hours after transduction, 200 μg/mL hygromycin was used to select for transduced cells and 1 μg/mL doxycycline was used to induce repression of the endogenous RUNX1. The screen was conducted with three biological replicates with greater than 1 million cells in each condition. At day 7 post transduction, the cells were split into two groups, 1 million cells for bulk RNA-seq and 100,000 cells for bulk ATAC-seq.

#### Single cell RNA sequencing library preparation

scRNA-seq experiments were performed with two replicates per condition (cells with and without doxycycline). Cells were first washed with a solution of PBS (Gibco) with 0.04% BSA (Gibco) by centrifuging the cells for 5 minutes at 300 G then resuspending them in the solution. After the wash, the cells were again centrifuged and resuspended in the same solution. The cells were filtered using a 40 μm cell strainer (VWR), and the concentration was determined using a manual hemacytometer (Thermo Fisher Scientific). The cells were then subjected to scRNA seq (10X genomics, chromium single cell 3’ v3, with two reactions per replicate) aiming for a target cell recovery of 10,000 cells per library. The single-cell libraries were generated according to manufacturer’s protocols with the following conditions: 11 PCR cycles run during cDNA amplification and 10 PCR cycles run during library generation. The libraries were sequenced using Illumina NovaSeq platform. To genotype the cells with the variant, the barcode sequences were amplified off of the cDNA pool generated in the scRNA-seq protocol. The barcodes were amplified via PCR using OneTaq 2X Master Mix (New England BioLabs) in 100 μL reactions, each split across 5 PCR tubes (20 μL per tube). For each sample the reactions contained 5 μL of primers RX1_07 and the NEBNext Universal PCR Primer for Illumina (New England BioLabs) (10 μM), 6 μL of cDNA, 50 μL of OneTaq, and the rest filled with water. A thermal cycler was used to heat the sample to 95 °C for 3 minutes, then 20 cycles of 98 °C for 20 seconds, 65 °C for 15 seconds, and 68 °C for 45 seconds, followed by a final 5 minute extension at 68 °C. The PCR products were purified using AMPure XP Beads beads at a 0.8:1 bead:PCR reaction ratio. The second step of PCR was performed. Subsequently, a NEBNext Ultra RNA Library Prep Kit (New England BioLabs) was used to generate Illumina compatible sequencing libraries; this was done in a 50 μL reaction split across 5 PCR tubes (10 μL per tube) with 20 ng of the first step purified PCR product.

### Single cell RNA sequencing library processing

#### Library fitness screening

A fitness screen was also performed concurrently with the single cell RNA sequencing screen. At days 2, 7, and 14 post-transfection, ∼1 million cells were collected, and their genomic DNA was isolated via a Qiagen DNeasy Blood and Tissue Kit. Barcodes corresponding to each library element at each timepoint, and replicate were then amplified from the genomic DNA using OneTaq 2X Master Mix. The sequencing libraries were amplified in 50 μL reactions, each split across 5 PCR tubes (10 μL per tube). For each sample, the reactions contained 2.5 μL of primers A and B (10 μM), 6 μg of gDNA, 25 μL of OneTaq, with the rest filled with water. The thermal cycler was used to heat the sample to 95 °C for 3 minutes, then 27 cycles of 98 °C for 20 seconds, 65 °C for 15 seconds, and 72 °C for 45 seconds, followed by a final 5 minute extension at 72 °C. The PCR products were purified using AMPure beads at 0.8:1 bead:PCR reaction ratio. NEBNext Multiplexed Oligos for Illumina (New England BioLabs) were then used to index the samples, and the samples were sequenced on an Illumina NovaSeq platform to a depth of 2.5 million reads/sample.

#### Freezing for bulk RNA-seq

Cells for bulk RNA-seq were pelleted and the media aspirated. They were flash-frozen in liquid nitrogen and stored at -80 °C.

#### Freezing for bulk ATAC-seq

Cells for bulk RNA-seq were pelleted in a centrifuge at 1000 G for 5 minutes at 4 °C, resuspended in cold PBS, and pelleted again. ATAC lysis buffer was made by mixing 100 μL 1M Tris-HCl pH 7.4, 20 μL 5 M NaCl, 30 μL 1M MgCl2, 100 μL 10% IGEPAL CA-630, and 9.75 mL water. The cells were lysed with the cold ATAC lysis buffer using 100 μL buffer per 100,000 cells and centrifuged at 1000 G for 10 minutes at 4 °C. The supernatant was removed, and the cells were flash-frozen in liquid nitrogen and stored at -80 °C.

#### Bulk RNA sequencing library preparation

Bulk RNA-seq experiments were performed with three replicates per condition. RNA was isolated from the cells using a Qiagen RNeasy Mini Kit according to the manufacturer’s protocols. Samples were prepared for bulk RNA-seq using the NEBNext Ultra II RNA Library Prep with Sample Purification Beads Kit (New England Biolabs) according to manufacturer’s protocols with the following conditions: 1 μg input RNA, library insert size = 200 nt. The bulk RNA-seq library was sequenced on an Illumina NovaSeq platform to a depth of 30 million reads/sample.

#### Bulk ATAC sequencing library preparation

Bulk ATAC-seq experiments were performed with three replicates per condition. Tagmentation buffer was prepared with 12.5 μL buffer, 9.75 μL H_2_O, 0.25 μL digitonin, and 2.5 μL Tn5 enzyme (Illumina) per sample. Each frozen cell pellet sample was resuspended in the tagmentation buffer and incubated at 37 °C for 45 minutes. 1x volume 40mM EDTA was added to each sample. The tagmented samples were purified using AMPure XP Beads at a 2:1 bead:tagmentation reaction ratio. The samples were incubated with the beads at room temperature for 15 minutes, then placed on a magnetic rack to separate the beads from the supernatant, which was discarded. The beads were washed twice with cold 80% ethanol, and the purified DNA was eluted from the beads using Buffer EB (Qiagen).

The tagmented DNA was dual indexed using i5 and i7 barcodes, giving each sample a unique barcode combination. The DNA and barcodes were added to NEB Hi Fidelity 2x PCR Mix (New England BioLabs) and amplified using the following PCR cycle: 72 °C for 7 minutes; 98 °C for 30 seconds; then 10 cycles of 98 °C for 10 seconds, 63 °C for 30 seconds, and 72 °C for 1 minute; and cooling back down to 4 °C. Double size selection was performed using AMPure XP Beads to select for the size of the final library. First, 0.55x volume AMPure Beads was added to each PCR reaction and incubated at room temperature for 15 minutes. The samples were placed on a magnetic rack and the supernatant transferred to new tubes, to which another 0.65x volume AMPure Beads were added (for a total of 1.2x volume PEG). The samples were incubated at room temperature for 15 minutes, the supernatant was discarded, and the beads were washed twice with cold 80% ethanol. DNA was eluted from the beads using Buffer EB and pooled together to make the final library for sequencing. The bulk ATAC-seq library was sequenced on an Illumina NovaSeq platform to a depth of 75 million reads/sample.

### Computational Analysis

#### scRNA-seq analysis

The single cell RNA sequencing screen was performed for two conditions: one treated with doxycycline to induce repression of the endogenous RUNX1 (named ‘dox’ condition), and the other not treated (named ‘nodox’ condition). The screening was conducted with two biological replicates for each condition, and single cell RNA sequencing was performed with two reactions per replicate, making a total of eight libraries: four containing cells treated with doxycycline (dox) and four not (nodox). Sequencing was run with a target cell recovery of 10,000 cells per library.

Sequencing reads in FASTQ format were aligned using the 10X Genomics Cell Ranger pipeline (version 3.1.0) [33], to the human transcriptome GRCh38 (version GRCh38-3.0.0), resulting in a gene by cell matrix of UMI counts for each library. To assign one or more genotypes to each cell, the plasmid barcode reads were aligned to GRCh38 using BWA, and labeled with its corresponding cell and UMI tags as described in the SEUSS pipeline [12] (https://github.com/yanwu2014/genotyping-matrices).

The UMI count matrices were processed using Seurat (version 4.1.0) [79]. Four dox and four nodox libraries were merged resulting in 86,120 cells and 21,153 genes, after removal of genes expressed in fewer than 3 cells. 44,418 cells not containing a genotype barcode, or containing more than one, were removed. To filter out low quality cells, we removed cells expressing fewer than 200 genes or more than 5000 genes. We also discarded cells that have over 20% of reads aligned to mitochondrial genes. Four perturbation variants (G138V, S145I, P157R, T161I) were excluded due to low cell counts (less than 10 cells for each condition), and one negative control was removed (G143G) due to a frame shift artifact during the mutation library preparation, resulting in 40,522 cells corresponding to 112 remaining variants (20,878 cells for dox, and 19,644 cells for nodox condition).

The count matrix was log-normalized with the default scale factor of 10,000 and the top 2000 variable genes were identified to be used for downstream analyses. Mitochondrial or ribosomal genes were not included in the top 2000 variable gene list. We then applied a linear transformation on the count matrix to center and scale the expression of each gene. We assigned cell cycle scores to each cell based on its expression of G2/M and S phase markers and applied a linear model to regress out effects of cell cycle heterogeneity. We performed linear dimensionality reduction (PCA) on the scaled data using the top 2000 variable genes (Supplementary Figure 2a).

#### T2 scores

In order to quantify the extent to which the expression profile of a variant deviates from the WT or LOF control variants, we used the Hotelling’s two-sample T-squared statistic (T2), a generalization of Student’s t-statistic that is used in a two-sample multivariate hypothesis testing [36]. For this comparison, we employed the principal component space, using the top 20 principal components (PC) to compare matrices of cells x 20 PCs for each variant. We used the hotellings2 function from the spm1d python package to compute the test statistics, named here as T2 scores. For each variant, first we compared against cells overexpressing the WT variant (T2 scores (vs. WT)), then we compared against cells overexpressing the LOF variant (T2 scores (vs. LOF)). Higher scores indicate a higher deviation from the variant being compared.

Based on T2 scores, for each variant, cells with the endogenous RUNX1 repressed (dox) displayed higher deviation from the WT or LOF control variants overall, in comparison to the cells carrying the endogenous RUNX1 (nodox) (**Supplementary Figure 2b-c**). Therefore, we decided to continue downstream analysis with dox condition cells only, corresponding to 20,878 cells with 20,389 genes. We repeated the previously described steps for log-normalization, identification of top 2000 variable genes, scaling, cell cycle effect regression, and PCA for these 20,878 dox cells.

#### Unsupervised clustering of single cells

Using the first 20 principal components, we clustered cells by first determining the nearest neighbors of each cell in the PCA space, and then by applying a modularity optimization algorithm that iteratively groups cells together with a resolution parameter of 0.3. We used UMAP, a non-linear dimensionality reduction technique [80], to visualize the three predicted unsupervised clusters where similar cells are placed together in low-dimensional space (**Figures 2a,b,f, Supplementary Figure 4c-d**). Unsupervised clusters were confirmed not to result from cell cycle phase heterogeneity, or batch effects from merging of four dox libraries (**Supplementary Figure 4c-d**).

We applied Fisher’s exact test to evaluate the enrichment or depletion of assigned phenotypes (**Figure 2g**), variant classes, or cell cycle phases (**Supplementary Figure 4a-b**) in each cluster. A log odds ratio (log(OR))>0 indicates enrichment, while a log(OR)<0 indicates depletion.

#### Unsupervised clustering of variants

For each variant, we computed the mean expression (log-normalized) of each of the top 2000 variable genes across all cells corresponding to the variant, resulting in an expression vector of size 2000 for each variant, representing its mean expression profile. This generated a count matrix of 112 variants by 2000 genes. We performed PCA on the count matrix, and using the first 20 principal components, we clustered variants by first determining the nearest neighbors of each variant in the PCA space, and then applying a modularity optimization algorithm that iteratively groups variants together with a resolution parameter of 0.8. We used UMAP to visualize the three predicted unsupervised clusters where similar variants are placed together in low-dimensional space (**Figures 2c-e**).

We performed differential gene expression analysis between variants using DESeq2 [81]. First, differentially expressed genes between WT and LOF control variants (203 genes; FDR<0.05) were obtained. Next, each hypomorphic variant was compared against the WT control variant separately, and genes that were differentially higher or lower expressed in at least one hypomorphic variant against WT (141 genes; FDR<0.05) were extracted. Then, the same procedure was performed against the LOF control variant (232 genes; FDR<0.05) (**Supplementary Figure 5a**).

#### Fitness analysis

To calculate fitness effects from genomic DNA reads, we first aligned reads to mutation barcodes (MagECK [82]) and counted the number of reads corresponding to each mutation for each replicate at each timepoint (days 2, 7 and 14 post transduction), resulting in a mutation by samples read counts matrix. We normalized read counts for each sample by dividing each column by its sum. We then divided read counts of each sample by the counts at day 2 post transduction, and log2 transformed it to obtain a measurement to represent fitness effects for each mutation and sample. We averaged fitness measurements from the two biological replicates taken at day 14 (**Figure 2i**) to compute mean fitness scores (**Figure 3d**).

#### Hierarchical variant clustering

We also hierarchically clustered variants based on Pearson correlation of their mean gene expression profiles using the top 2000 variable genes. We ordered the leaves of the resulting dendrogram by increasing T2 scores obtained from comparison to the WT variant. To obtain discrete cluster assignments, we cut the hierarchy based on visual inspection, obtaining three main variant clusters that largely agree with WT-like, hypomorphic and LOF-like annotations (only 1 variant difference for the hypomorphic/LOF-like separation). We further cut the hierarchy of the middle cluster, representing hypomorphic variants, into three sub-clusters: named as hypomorphic-1, hypomorphic-2 and hypomorphic-3.

#### Hierarchical gene clustering

To determine genes whose expression is impacted by variants, we hierarchically clustered genes based on Manhattan distance between them using mean gene expression profiles of variants, resulting in gene groups with various expression profiles across variant clusters.

#### Cell state analysis

We applied a non-negative matrix factorization algorithm (CoGAPS [52] on the expression matrix of the top 2000 variable genes of 14,217 cells harboring perturbation variants or WT or LOF control constructs, using default parameters, which produced a gene by pattern (2000 x 7) and a pattern by cell (7 x 14,217) matrix. Using the pattern by cell matrix, we hierarchically clustered the top 2000 variable cells into 7 clusters which roughly corresponds to the 7 identified patterns (**Supplementary Figure 7a**), and applied a Fisher’s exact test to evaluate the enrichment or depletion of variant phenotypic annotations for each cluster (**Supplementary Figure 7b**). Using the gene by pattern matrix, we assigned non-overlapping gene markers for each pattern by distributing genes into patterns with the lowest ranking.

#### Predicting variant transcriptomic effects

To generate a binary classification task, we divided the 79 RUNX1-perturbing variants in our library into a positive (38 functional variants: 24 LOF-like and 14 hypomorphic) and negative class (41 WT-like variants). First, we obtained 85 features for each variant from the SNVBox database [58] describing substitution effects on amino acid biophysical properties, evolutionary conservation of variant sites, local sequence biases, and site-specific functional annotations. Then, we performed a 60-40 random split on the dataset, to generate a training (n=49: 24 positive, 25 negative) and a test set (n=30: 14 positive, 16 negative) with balanced ratios of positive and negative class variants.

We trained a Random Forest classifier (n_estimators = 1000, max_features = ’sqrt’) on the training set using the scikit-learn Python package and tested on the test set. The classifier score (between 0-1) represents the percentage of decision trees that classify a mutation as positive. Receiver Operator Characteristic (ROC) and Precision–Recall curves (PR) were constructed from the classifier scores and the AUC statistic was used as a measure of classifier performance.

Next, we trained another Random Forest classifier using the entire dataset of 79 perturbation variants and predicted transcriptomic effect labels for all remaining possible missense mutations on the RUNX1 protein (n=2594). We used the positive (5 core mutations) and negative (7 predicted neutral mutations) missense control variants in our RUNX1 mutation library as a validation set.

#### MLL dataset

Patient samples sent to the Munich Leukemia Laboratory (MLL) for routine diagnostic workup between August 2005 and March 2023 and that were diagnosed with AML were queried for missense mutations in RUNX1. AML diagnoses were based on cytomorphology, immunophenotype, cytogenetics, and molecular genetics following gold standard practices. All patients gave their written informed consent for scientific evaluations. The study was approved by the Internal Review Board and adhered to the tenets of the Declaration of Helsinki. In total, 716 individuals from the MLL cohort carried 1 or more missense mutations in the Runt domain (amino acid positions 50-177), totalling 772 mutations. Mutations were defined with respect to the ENST00000344691 transcript of RUNX1.

#### Selecting variants for validation with bulk sequencing

Ten hypomorphic variants showing the largest deviation from control conditions based on mean of T2_WT_ and T2_LOF_ scores were selected: two hypomorphic-I (N82I, P156R), seven hypomorphic-II (L62P, L94P, G95R, V97D, G100V, V137D, I166S) and one hypomorphic-III (R118G). L94P was removed for being predicted to target similar protein interactions as G95R, and a LOF-like variant (V159D) predicted to be involved in RUNX1-CBFB binding was added. WT and LOF control variants were included bringing the total number of variants chosen for bulk sequencing to 12.

#### Bulk RNA-seq analysis

Sequencing reads in FASTQ format were aligned to the human transcriptome GRCh38 (Gencode v30 - GRCh38.p12) using STAR (version 2.7.1a) [83]. RSEM (version 1.3.1) is used to calculate read counts for each sample and replicate (‘rsem-calculate-expression’ command), and to generate a gene by sample matrix (‘rsem_generate_data_matrix’ command) of the raw counts (‘expected_count’ column). Starting with 57,535 gene features, we removed genes with less than 10 reads in total across all the samples, along with mitochondrial and ribosomal genes, resulting in 18,646 remaining genes.

We first normalized raw counts using the variance stabilizing transformation, which transforms counts on the log2 scale and normalizes with respect to library size [81]. We removed two outlier samples (replicates 2 of samples with N82I and V137D mutations) identified based on expression profiles of top 2000 variable genes (**Supplementary Figure 11**) and removed batch effects between replicates using the limma package [84]. For visualization purposes, we averaged gene expression across replicates for each sample. In order to validate variant clustering results obtained from scRNA-seq here in the bulk setting, we used top 2000 variable genes obtained from the scRNA-seq analysis, to perform PCA and hierarchical clustering of samples and genes based on Manhattan distance. We ordered the leaves of the resulting sample dendrogram by increasing T2 scores obtained from the scRNA-seq analysis by comparison to the WT variant. The same analysis was also performed using all replicates instead of their means (**Supplementary Figure 9**).

Differential expression analysis between samples was performed with DESeq2 [81]. Each hypomorphic variant was compared against the WT and LOF control variants separately, and genes that were differentially (FDR<0.05) upregulated (202 genes) or downregulated (89 genes) in at least one hypomorphic variant against both WT and LOF controls were extracted.

RNA expression coverage tracks (bigWig files) were generated from BAM format using deepTools bamCoverage [85].

#### Bulk ATAC-seq analysis

Sequencing reads in FASTQ format were aligned to the human transcriptome GRCh38 and processed using the nf-core/atacseq pipeline (version 1.2.2) [86], built using Nextflow (version 22.04.0), in conjunction with Singularity. The command used is ‘nextflow run nf-core/atacseq -r master -name “run/name” -profile “singularity” -work-dir “work/directory/path” -params-file “params/file/path” --genome GRCh38 --narrow_peak true’, with default parameters. First, fastq files from two ATAC-seq runs were merged with “cat” command for each read (reads 1 and 2 for paired-end data) of each replicate (three biological replicates) of each sample (12 samples); and the pipeline was run on the merged FASTQ files. Briefly, the pipeline performs adapter trimming using Trim Galore! (https://www.bioinformatics.babraham.ac.uk/projects/trim_galore/), read alignment with BWA [87], filtering with SAMtools [88] (e.g. removal of mitochondrial reads), BEDTools [89], BamTools [90], Pysam (https://github.com/pysam-developers/pysam), and picard (https://broadinstitute.github.io/picard/), normalized coverage track generation with BEDTools and bedGraphToBigWig [91], genome-wide enrichment with deepTools [85], peak calling with MACS2 [92] (narrow peaks), and quality control and statistics reporting with MultiQC [93].

Coverage tracks were further processed with AtacWorks [94], which uses a deep learning model trained on high quality ATAC-seq data to remove background noise. ATAC-seq data from K562 cells was obtained from the ENCODE data portal [95] (experiment ENCSR868FGK). To generate a model of noisy data, the aligned reads files from replicate 1 (ENCFF534DCE) were subsampled to about 20 million reads with SAMtools view [88], and converted to bigWig format with deepTools bamCoverage [85]. The resulting bigWig file and the bigWig for the entirety of replicate 1 (ENCFF670QXU) were provided to AtacWorks as the noisy versus clean data to train the model.

The two highest quality replicates for each RUNX1 variant (bigWig files) were denoised using the trained AtacWorks model and combined with UCSC bigWigMerge [91]. Peaks were called on the summed files using MACS2 callpeak [92] and were compared against the ENCODE K562 ATAC data and filtered as follows.

For the wild type sample:

1. denoised peaks that were not observed in the undenoised peak set <u>or</u> in the ENCODE peak set were marked as noise and removed,
2. peaks that were seen in the denoised bigWig track but lost during peak calling <u>and</u> that were also observed in either the undenoised or the ENCODE peak set were rescued.

For the other samples:

3. denoised peaks that were not observed in the undenoised peak set <u>or</u> in the ENCODE peak set were marked as noise and removed,
4. peaks that were seen in the denoised bigWig track but lost during peak calling <u>and</u> that were also observed in the ENCODE peak set were rescued as “wild type” peaks,
5. peaks that were seen in the denoised bigWig track but lost during peak calling <u>and</u> that were also observed in the undenoised peak set were rescued as “mutation” peaks.

The filtered peaks were merged with BEDTools merge [89] to generate the final denoised consensus peak set. On the consensus peak set, read counts were obtained with featureCounts [96]. Using HOMER [97], enriched motifs were identified (findMotifsGenome function) and filtered for Runt domain motifs (5 motifs total) to represent RUNX1 DNA binding sites. Consensus peaks were annotated with genomic features and Runt motifs (within 1000 base pairs) using HOMER (annotatePeaks function). They were also annotated with ChromHMM states [98] using a 25-state model for K562 cells obtained from the Roadmap Epigenomics Project (https://egg2.wustl.edu/roadmap/web_portal/). Peaks that overlap with Hi-C loops [99] were identified with BEDTools pairtobed [89] using Hi-C chromatin loop data for K562 cells obtained from NCBI GEO (GSM1551620).

We identified peaks as promoters if located within 1 kbp downstream and 100 bp upstream of the transcription start site (TSS) of a gene. We identified peaks as enhancers if annotated with a ChromHMM enhancer state. We filtered both promoter and enhancer peaks that contain Runt motifs, to study genes regulated by RUNX1. Genes associated with each enhancer peak were identified using the Hi-C chromatin loops.

RNA expression and DNA accessibility coverage tracks, ChromHMM states and Hi-C loops were visualized using CoolBox [100].

#### Statistical analysis

Correlations were evaluated using the Pearson correlation coefficient. Odds ratios were calculated using Fisher’s exact test. Distributions were compared using a Mann–Whitney U test. Multiple testing correction is applied where applicable using the Benjamini-Hochberg method.

Gene set overrepresentation analysis is performed using the Gene Ontology (GO) biological process terms (2021) or Reactome pathways (2022), with the EnrichR package [101].

#### Data and code availability

An SRA accession for all sequencing data is pending and code is available under an MIT license via Github repository https://github.com/cartercompbio/Runt_domain_muts.

