## Supplementary materials for "Interface-guided phenotyping of coding variants in the transcription factor RUNX1 with SEUSS"

### SUPPLEMENTARY FIGURES

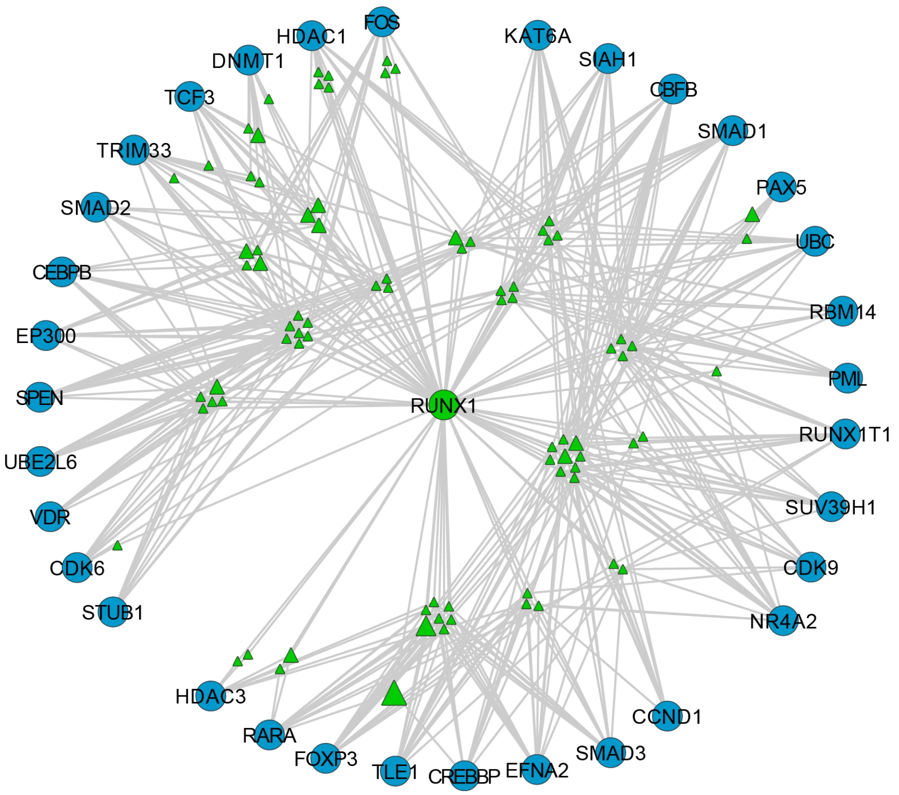


**Supplementary Figure 1. RUNX1 interaction network.**
RUNX1 protein (green circle), its interaction partners (blue circles), and the interface residues of RUNX1 by which it physically interacts with each partner (green triangles) are displayed. Triangle size represents the number of human tumors in which the residue was mutated.

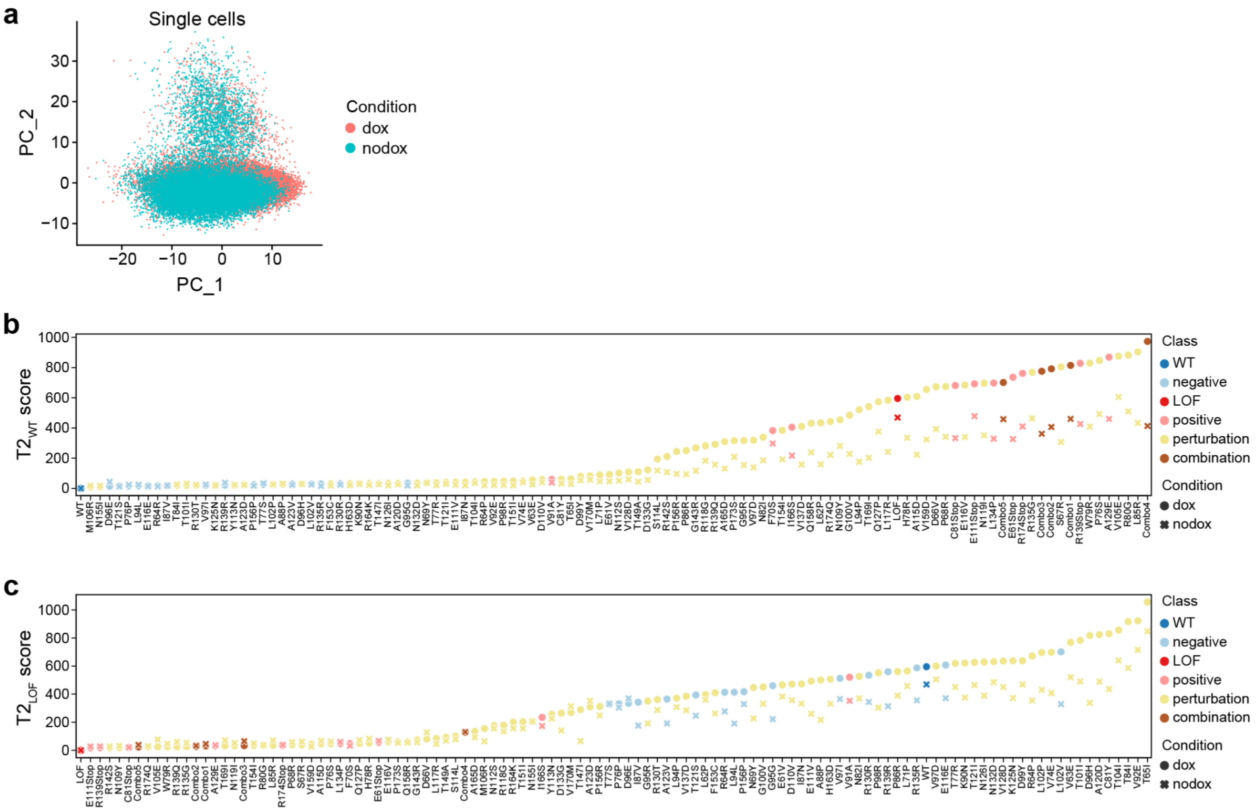


**Supplementary Figure 2. Comparison of RUNX1 variant transcriptional effects between cells treated with doxycycline (dox) or not (nodox).**

**a.** PCA plot of single cells colored by doxycycline treatment condition, obtained using the top 2000 variable genes. Cell cycle effects are regressed out.

**b-c.** T2 scores of each variant for cells with dox (circle) or nodox (cross) condition, when compared against **b.** the WT or **c.** LOF control, colored by variant classes. Higher scores indicate a higher deviation from the control variant being compared. Variants are ordered by increasing T2 scores for the dox condition.

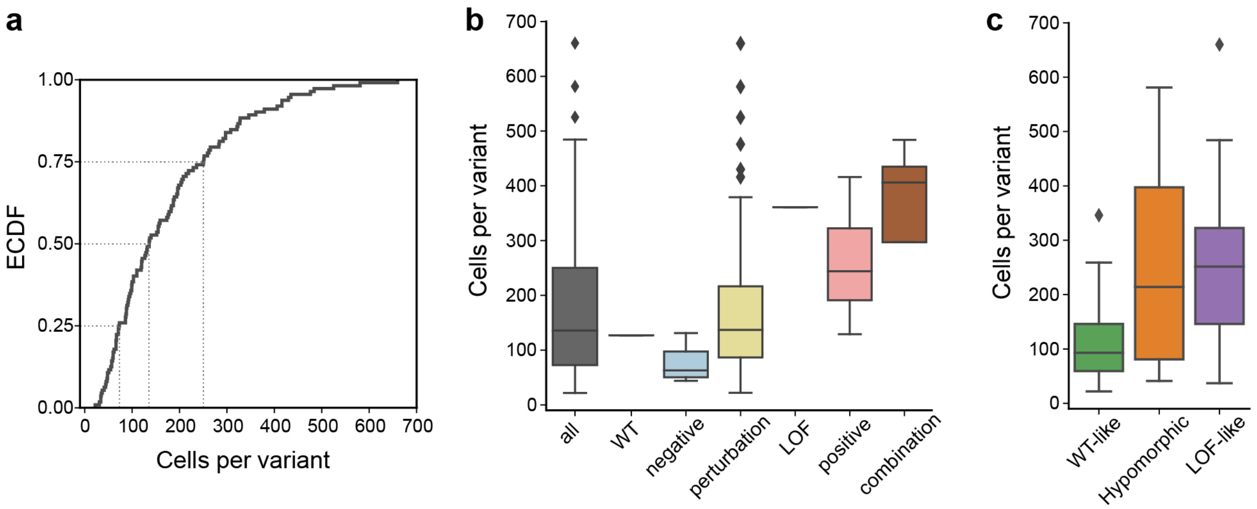


**Supplementary Figure 3. Distribution of number of cells per variant.**

**a.** Empirical cumulative distribution function (ECDF) of number of cells profiled for each variant (median 136 cells per variant).

**b-c.** Distribution of number of cells per variant for **b.** each variant class, or **c.** for each variant assigned phenotype.

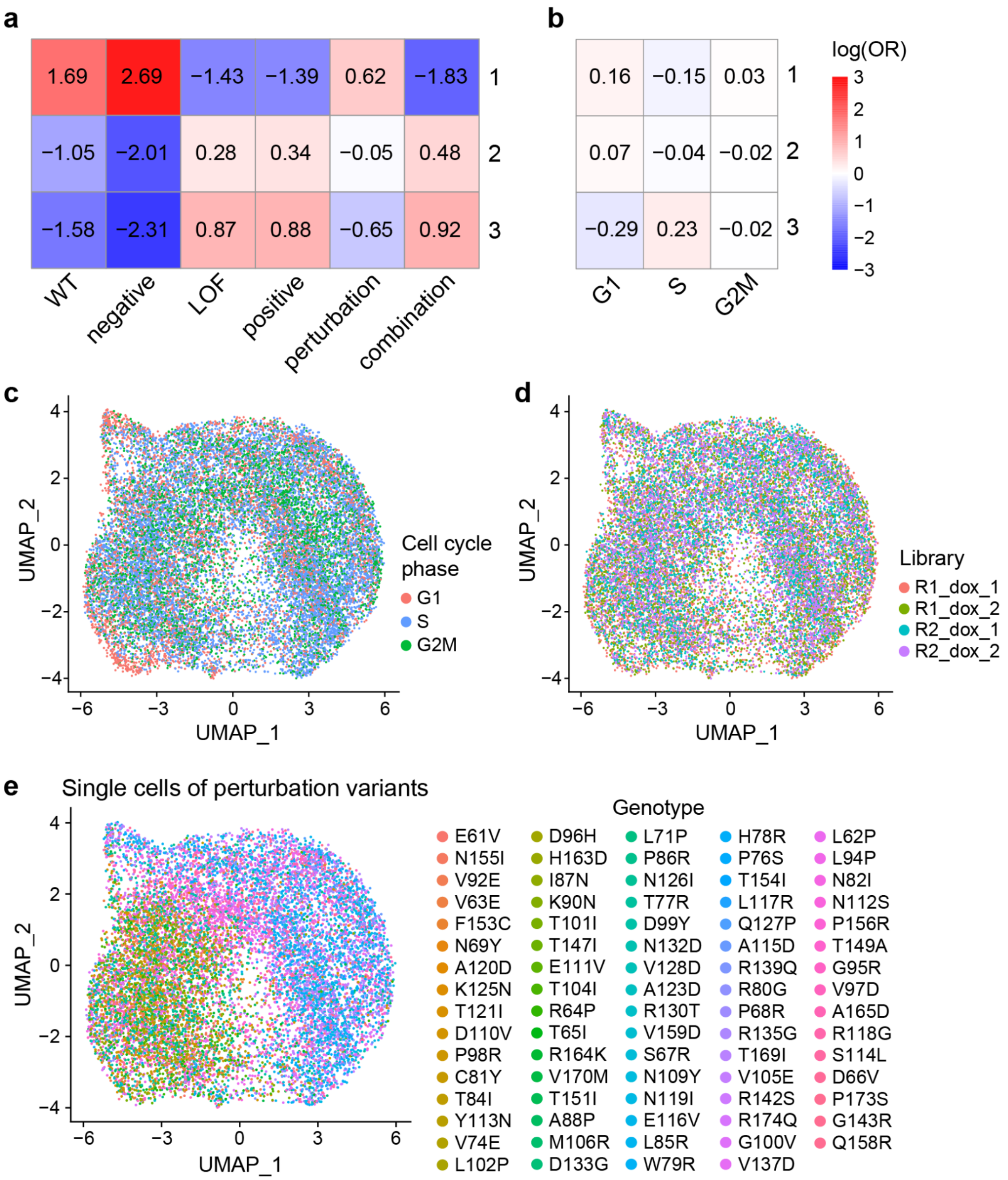


**Supplementary Figure 4. Unsupervised analysis of RUNX1 variant transcriptional effects.**

**a-b.** Cluster enrichment of single cells (unsupervised clusters from Figure 2a) for **a.** variant classes (Figure 2b), and **b.** cell cycle phases, based on log of odds ratios obtained using Fisher's exact test. Positive values indicate enrichment, while negative values indicate depletion.

**c-d.** UMAP embedding of single cells carrying any of the 112 library variants, colored by **c.** cell cycle phases, or **d.** dox libraries, obtained using the top 2000 variable genes. Cell cycle effects are regressed out.

**e.** UMAP embedding of single cells containing perturbation variants only, colored by genotypes, obtained using the top 2000 variable genes. Cell cycle effects are regressed out.

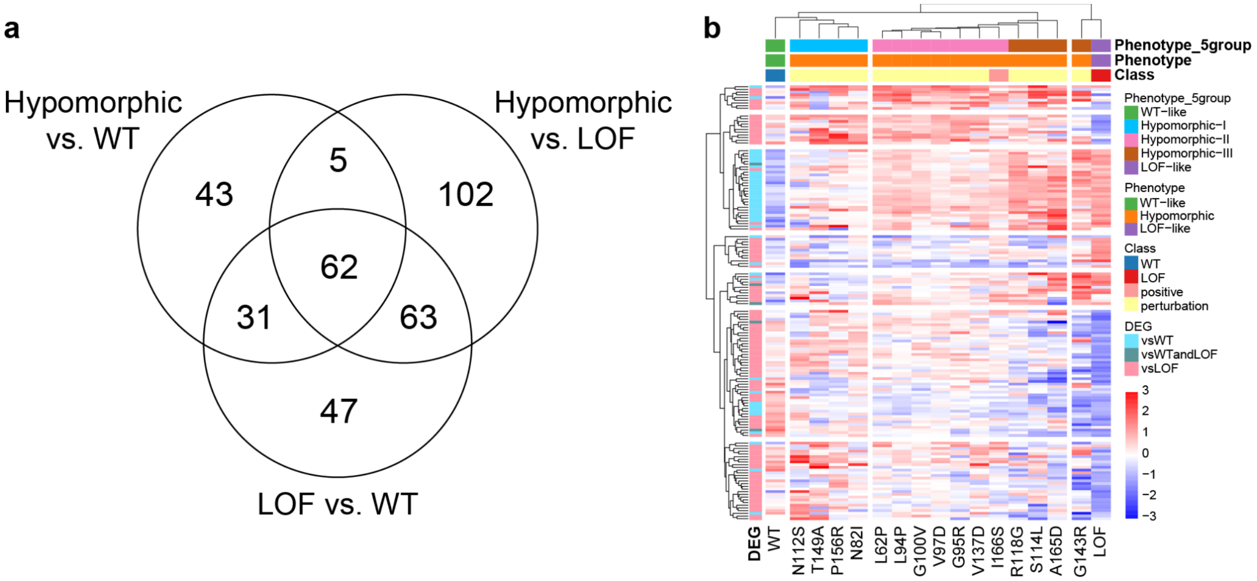


**Supplementary Figure 5. Differential expression of genes in hypomorphic variants against WT and LOF controls.**

**a.** Venn diagram displaying the number of genes that are differentially expressed between single cells harboring WT vs. LOF control variants, a hypomorphic variant vs. WT, or a hypomorphic variant vs. LOF control variant.

**b.** Heatmap showing mean expression profiles of 150 genes (rows) that are differentially higher or lower expressed in a hypomorphic variant against the WT (light blue) or LOF (pink) control variant (columns), or both (teal), but not between WT vs. LOF controls. Genes and variants are hierarchically clustered into seven and four clusters, respectively. The leaves of the variant dendrogram are ordered by increasing $T2_{WT}$ scores. Gene expression values are z-scored.

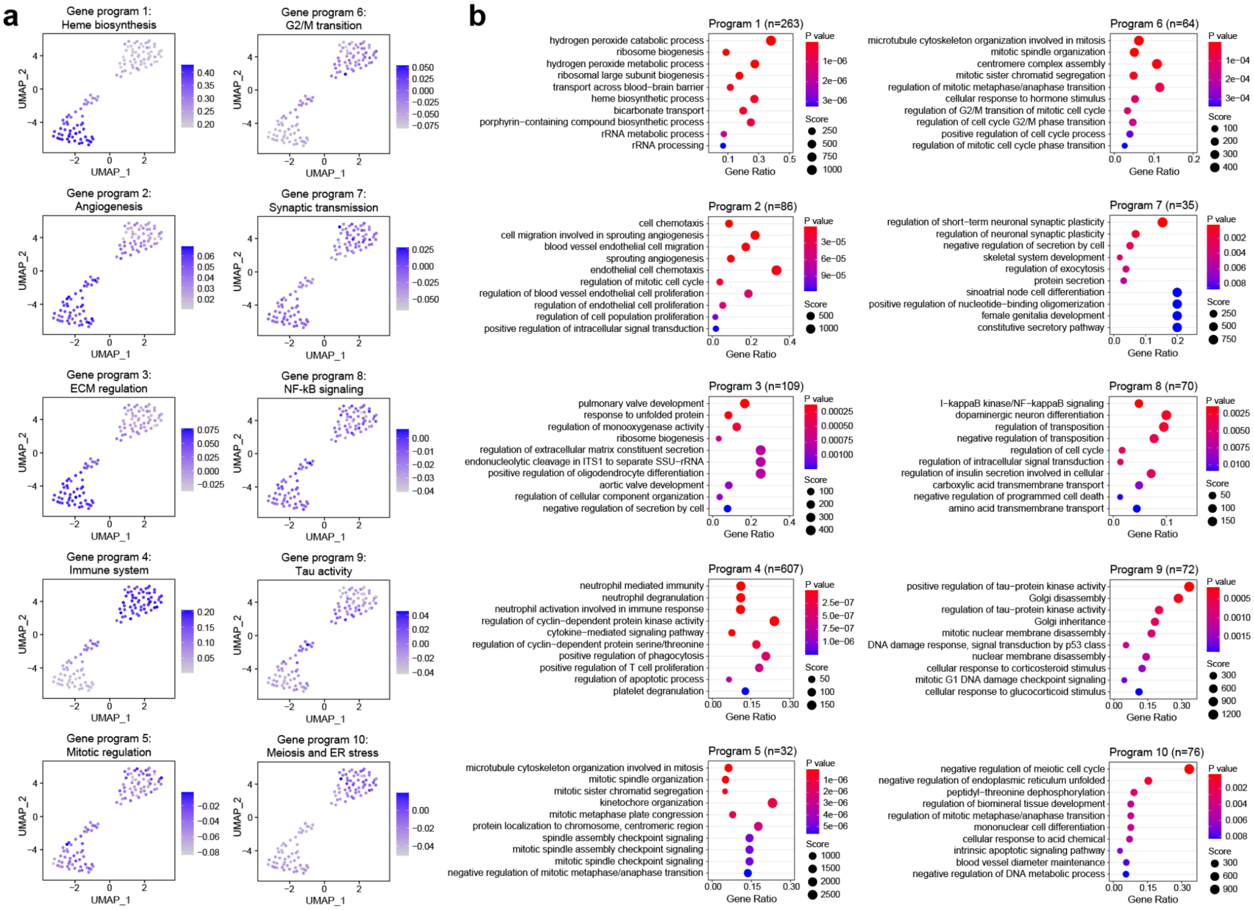


**Supplementary Figure 6. Hierarchical clustering based gene expression programs.**

**a.** Aggregated mean expression of genes for each gene program (Figure 3a) across cells for each variant. **b.** Gene set overrepresentation analysis results for GO Biological Process terms for each gene program (Figure 3a) displaying top 10 terms.

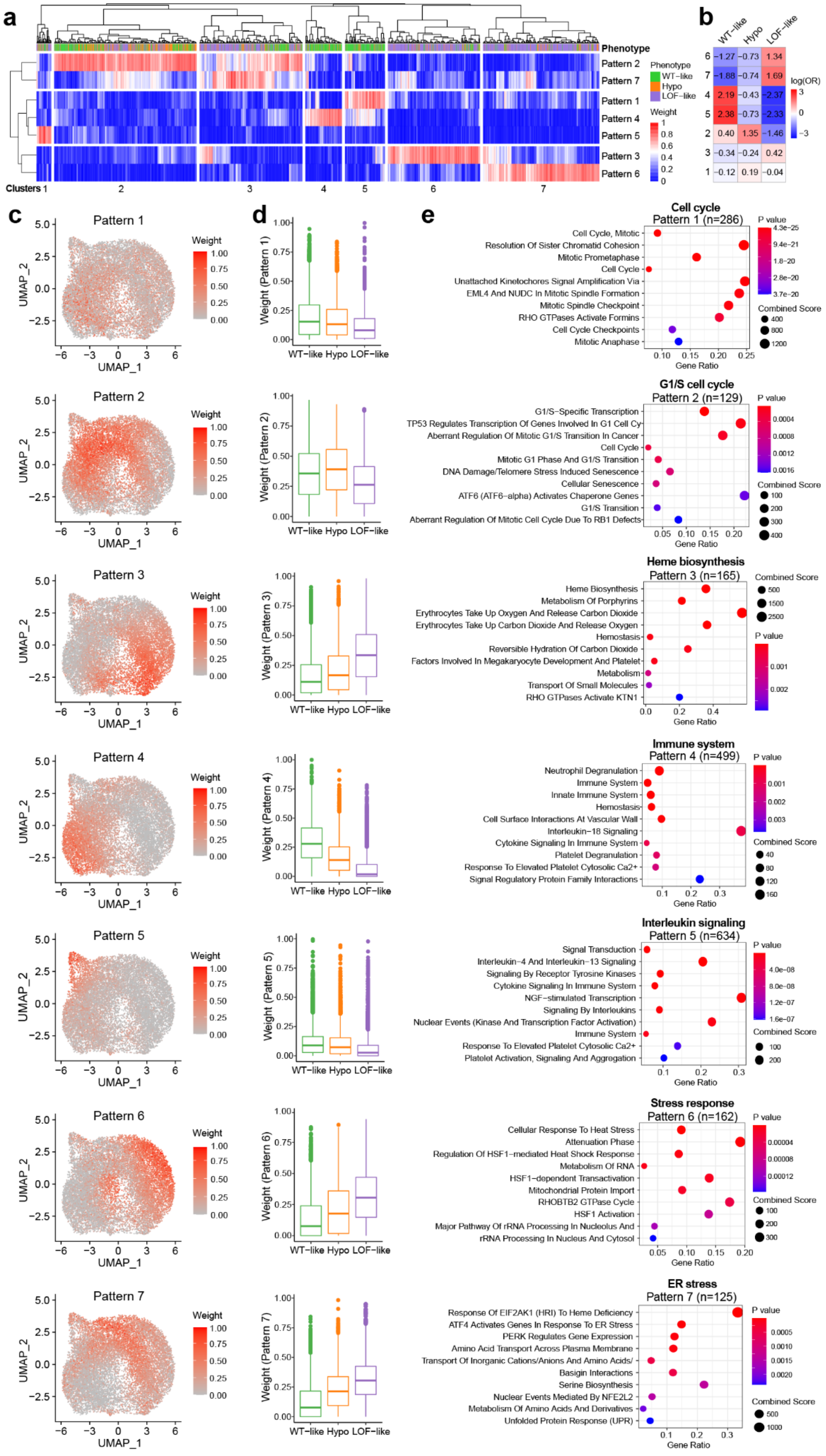
a
Phenotype
Pattern 2
Pattern 7
Pattern 1
Pattern 4
Pattern 5
Pattern 3
Pattern 6
Phenotype
WT−like
Hypo
LOF−like
Weight
Clusters 1
2
3
4
5
6
7
b
WT−like
Hypo
LOF−like
6 −1.27 −0.73 1.34
7 −1.88 −0.74 1.69
4 2.19 −0.43 −2.37
5 2.38 −0.73 −2.33
2 0.40 1.35 −1.46
3 −0.34 −0.24 0.42
1 −0.12 0.19 −0.04
log(OR)
c
Pattern 1
Pattern 2
Pattern 3
Pattern 4
Pattern 5
Pattern 6
Pattern 7
UMAP_1
UMAP_2
Weight
d
Weight (Pattern 1)
Weight (Pattern 2)
Weight (Pattern 3)
Weight (Pattern 4)
Weight (Pattern 5)
Weight (Pattern 6)
Weight (Pattern 7)
WT−like Hypo LOF−like
e
Cell cycle
Pattern 1 (n=286)
Cell Cycle, Mitotic
Resolution Of Sister Chromatid Cohesion
Mitotic Prometaphase
Cell Cycle
Unattached Kinetochores Signal Amplification Via
EML4 And NUDC In Mitotic Spindle Formation
Mitotic Spindle Checkpoint
RHO GTPases Activate Formins
Cell Cycle Checkpoints
Mitotic Anaphase
G1/S cell cycle
Pattern 2 (n=129)
G1/S−Specific Transcription
TP53 Regulates Transcription Of Genes Involved In G1 Cell Cy
Aberrant Regulation Of Mitotic G1/S Transition In Cancer
Cell Cycle
Mitotic G1 Phase And G1/S Transition
DNA Damage/Telomere Stress Induced Senescence
Cellular Senescence
ATF6 (ATF6−alpha) Activates Chaperone Genes
G1/S Transition
Aberrant Regulation Of Mitotic Cell Cycle Due To RB1 Defects
Heme biosynthesis
Pattern 3 (n=165)
Heme Biosynthesis
Metabolism Of Porphyrins
Erythrocytes Take Up Oxygen And Release Carbon Dioxide
Erythrocytes Take Up Carbon Dioxide And Release Oxygen
Hemostasis
Reversible Hydration Of Carbon Dioxide
Factors Involved In Megakaryocyte Development And Platelet
Metabolism
Transport Of Small Molecules
RHO GTPases Activate KTN1
Immune system
Pattern 4 (n=499)
Neutrophil Degranulation
Immune System
Innate Immune System
Hemostasis
Cell Surface Interactions At Vascular Wall
Interleukin−18 Signaling
Cytokine Signaling In Immune System
Platelet Degranulation
Response To Elevated Platelet Cytosolic Ca2+
Signal Regulatory Protein Family Interactions
Interleukin signaling
Pattern 5 (n=634)
Signal Transduction
Interleukin−4 And Interleukin−13 Signaling
Signaling By Receptor Tyrosine Kinases
Cytokine Signaling In Immune System
NGF−stimulated Transcription
Signaling By Interleukins
Nuclear Events (Kinase And Transcription Factor Activation)
Immune System
Response To Elevated Platelet Cytosolic Ca2+
Platelet Activation, Signaling And Aggregation
Stress response
Pattern 6 (n=162)
Cellular Response To Heat Stress
Attenuation Phase
Regulation Of HSF1−mediated Heat Shock Response
Metabolism Of RNA
HSF1−dependent Transactivation
Mitochondrial Protein Import
RHOBTB2 GTPase Cycle
HSF1 Activation
Major Pathway Of rRNA Processing In Nucleolus And
rRNA Processing In Nucleus And Cytosol
ER stress
Pattern 7 (n=125)
Response Of EIF2AK1 (HRI) To Heme Deficiency
ATF4 Activates Genes In Response To ER Stress
PERK Regulates Gene Expression
Amino Acid Transport Across Plasma Membrane
Transport Of Inorganic Cations/Anions And Amino Acids/
Basigin Interactions
Serine Biosynthesis
Nuclear Events Mediated By NFE2L2
Metabolism Of Amino Acids And Derivatives
Unfolded Protein Response (UPR)
Gene Ratio
P value
Combined Score

**Supplementary Figure 7. Gene expression patterns identifying cell states for WT-like, hypomorphic or LOF-like variants.**

**a.** Heatmap showing pattern weights of single cells (columns) for each of 7 patterns (rows) identified by non-negative matrix factorization. Cells are clustered into seven clusters which roughly correspond to the 7 patterns. Cells are colored by phenotype of the variant they harbor.

**b.** Cluster enrichment of single cells (hierarchical clusters from **a**) for variant phenotypes based on log of odds ratios obtained using Fisher's exact test. Positive values indicate enrichment, while negative values indicate depletion.

**c.** UMAP embedding of single cells, colored by pattern weights for each of 7 patterns.

**d.** Boxplots of pattern weights of single cells, across variant phenotypes.

**e.** Gene set overrepresentation analysis of marker genes of each of 7 patterns for Reactome pathways. Top 10 terms ordered by p-values are displayed.

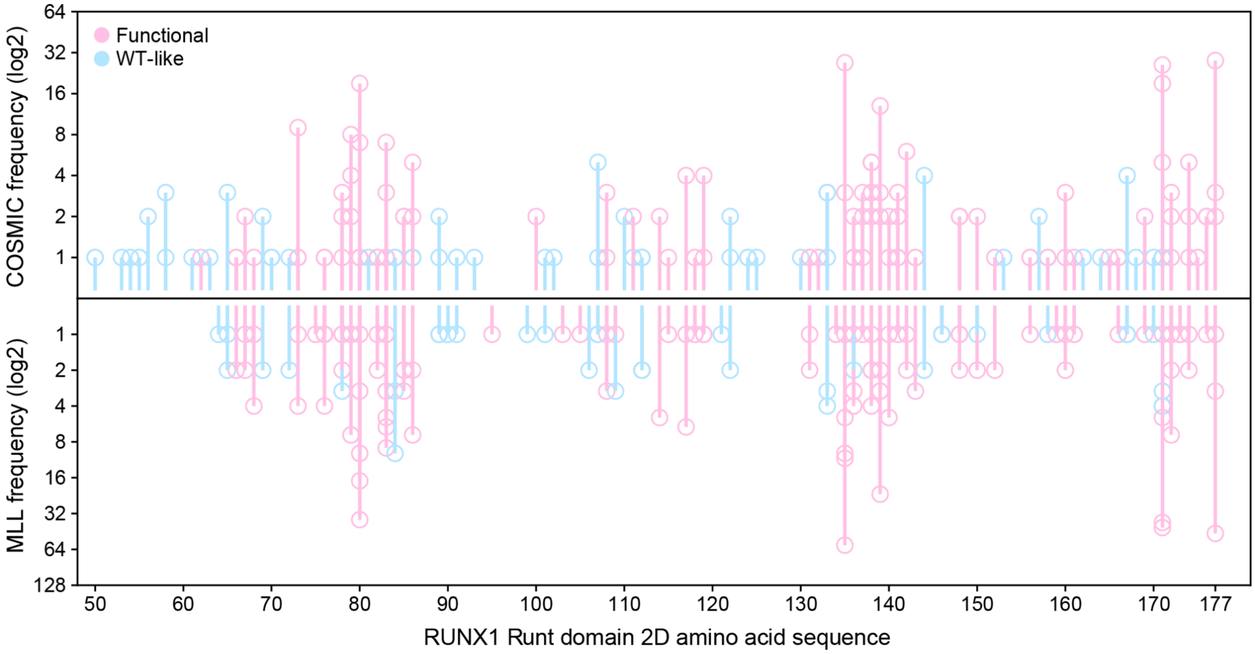


**Supplementary Figure 8. Sequence-based phenotypic profiling of RUNX1 Runt domain mutation missense mutations observed in human tumors (Cosmic or MLL cohorts).**
Frequency of mutations in COSMIC (top panel), or MLL (bottom panel) cohorts (log2 scaled), distributed across 2D amino acid sequence of RUNX1 Runt domain. Mutations are colored by transcriptomic effect labels predicted by our RUNX1-based model (pink: functional, blue: WT-like). Mutations overlapping with our RUNX1 variant library are excluded.

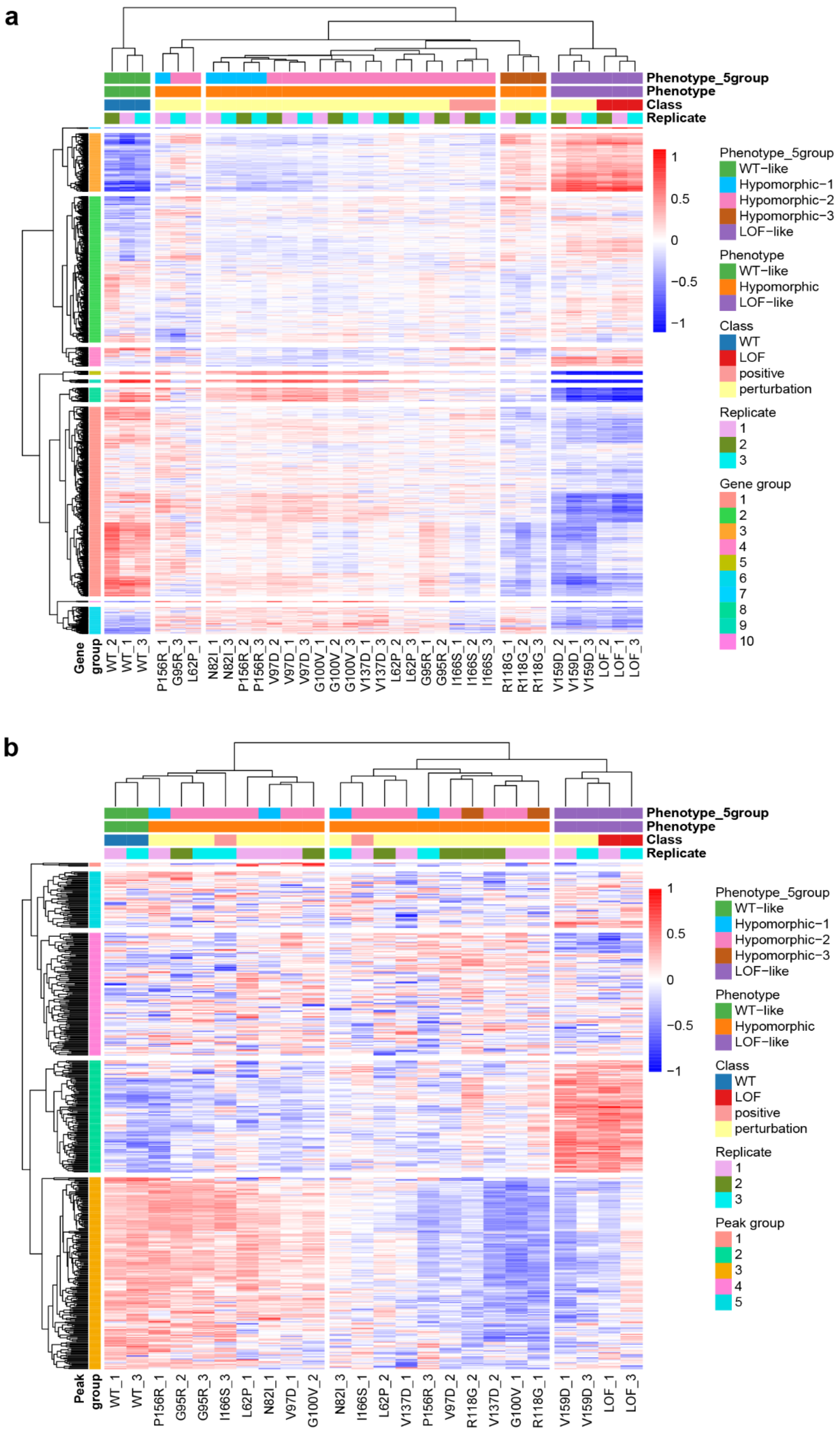
a
Phenotype_5group
Phenotype
Class
Replicate
Phenotype_5group
WT−like
Hypomorphic−1
Hypomorphic−2
Hypomorphic−3
LOF−like
Phenotype
WT−like
Hypomorphic
LOF−like
Class
WT
LOF
positive
perturbation
Replicate
1
2
3
Gene group
1
2
3
4
5
6
7
8
9
10
1
0.5
0
−0.5
−1
Gene group
WT_2
WT_1
WT_3
P156R_1
G95R_3
L62P_1
N82I_1
N82I_3
P156R_2
P156R_3
V97D_2
V97D_1
V97D_3
G100V_1
G100V_2
G100V_3
V137D_1
V137D_3
L62P_2
L62P_3
G95R_1
G95R_2
I166S_1
I166S_2
I166S_3
R118G_1
R118G_2
R118G_3
V159D_2
V159D_1
V159D_3
LOF_2
LOF_1
LOF_3
b
Phenotype_5group
Phenotype
Class
Replicate
Phenotype_5group
WT−like
Hypomorphic−1
Hypomorphic−2
Hypomorphic−3
LOF−like
Phenotype
WT−like
Hypomorphic
LOF−like
Class
WT
LOF
positive
perturbation
Replicate
1
2
3
Peak group
1
2
3
4
5
1
0.5
0
−0.5
−1
Peak group
WT_1
WT_3
P156R_1
G95R_2
G95R_3
I166S_3
L62P_1
N82I_1
V97D_1
G100V_2
N82I_3
I166S_1
L62P_2
V137D_1
P156R_3
V97D_2
R118G_2
V137D_2
G100V_1
R118G_1
V159D_1
V159D_3
LOF_1
LOF_3

**Supplementary Figure 9. Bulk RNA- and ATAC-seq analysis of 12 validation variants with all replicates, related to Figure 5.**

**a.** Hierarchical clustering of samples (rows) and genes (columns) in the bulk RNA-seq setting, using top 2000 variable genes obtained from scRNA-seq. All sample replicates are present. The leaves of the variant dendrogram are ordered by increasing $T2_{WT}$ scores. Gene expression values are z-scored.

**b.** Hierarchical clustering of samples (rows) and peaks (columns) in the bulk ATAC-seq setting, using top 500 variable peaks. The two highest quality replicates of each sample are present. The leaves of the variant dendrogram are ordered by increasing $T2_{WT}$ scores. DNA accessibility values are z-scored.

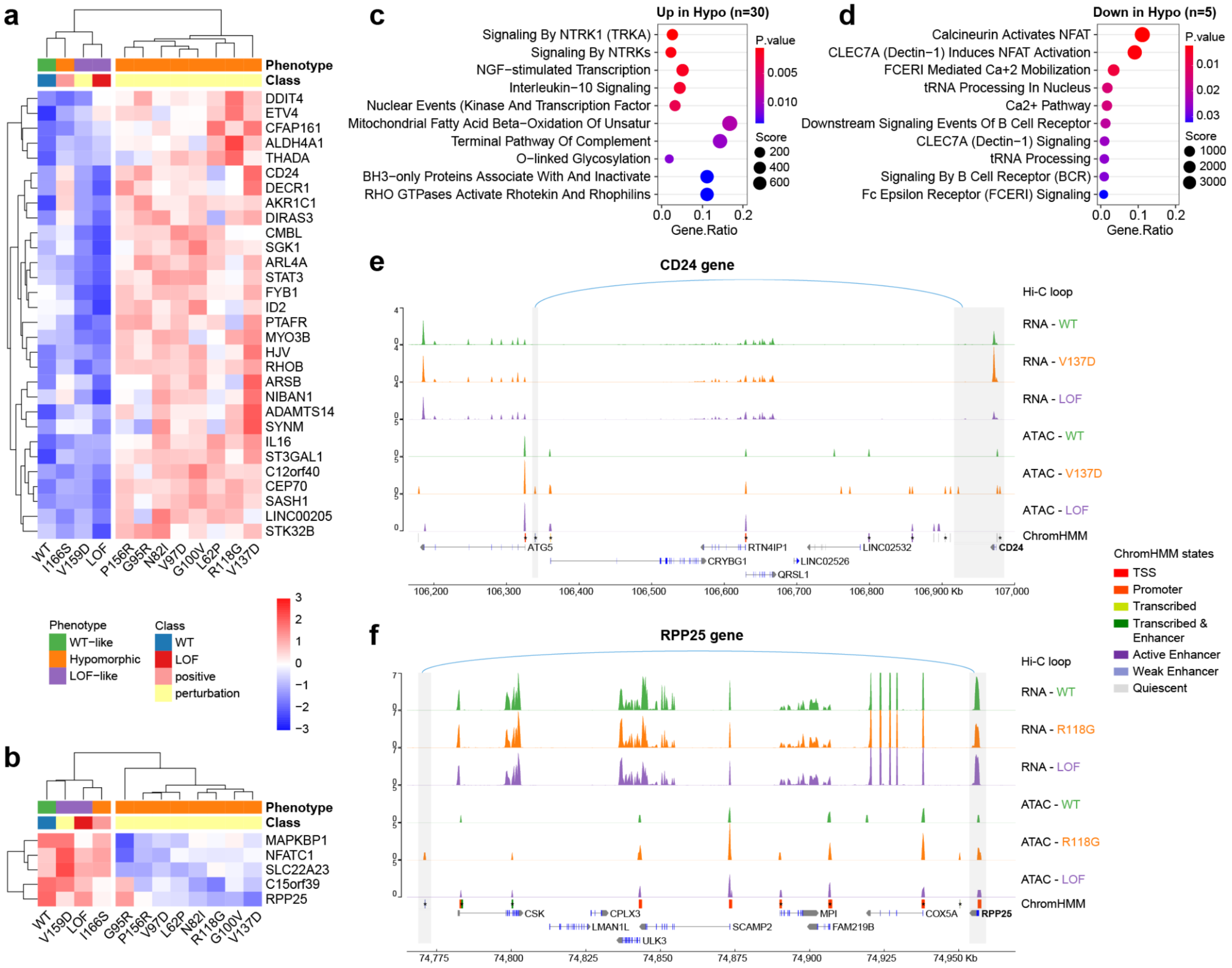


**Supplementary Figure 10. Regulatory consequences on RUNX1 binding at enhancer regions for hypomorphic Runt domain variants.**

**a-b.** Hierarchical clustering of samples (rows) and genes (columns) in the bulk RNA-seq setting, for genes that have a nearby enhancer with a RUNX1 binding site and are significantly **a.** upregulated (n=30), or **b.** downregulated (n=5), in at least one hypomorphic variant against both WT and LOF controls. Gene expression is averaged across replicates. Gene expression values are z-scored.

**c-d.** Gene set overrepresentation analysis results for Reactome pathways for up- and down-regulated genes. Top 10 terms ordered by p-values are displayed.

**e.** RNA-seq, ATAC-seq and Hi-C tracks illustrate an enhancer peak linked to the CD24 promoter in the context of the V137D variant, potentially explaining why this gene is expressed more highly in the hypomorphic context.

**f.** RNA-seq, ATAC-seq and Hi-C tracks link an enhancer peak 5' to CSK to decreased expression of RPP25 in the R118G variant but not other genes in the region. ATAC-seq peaks are annotated with ChromHMM states, and star (*) indicates the presence of a RUNX motif. Exons and UTRs of each gene are displayed with blue and gray bands, accordingly.

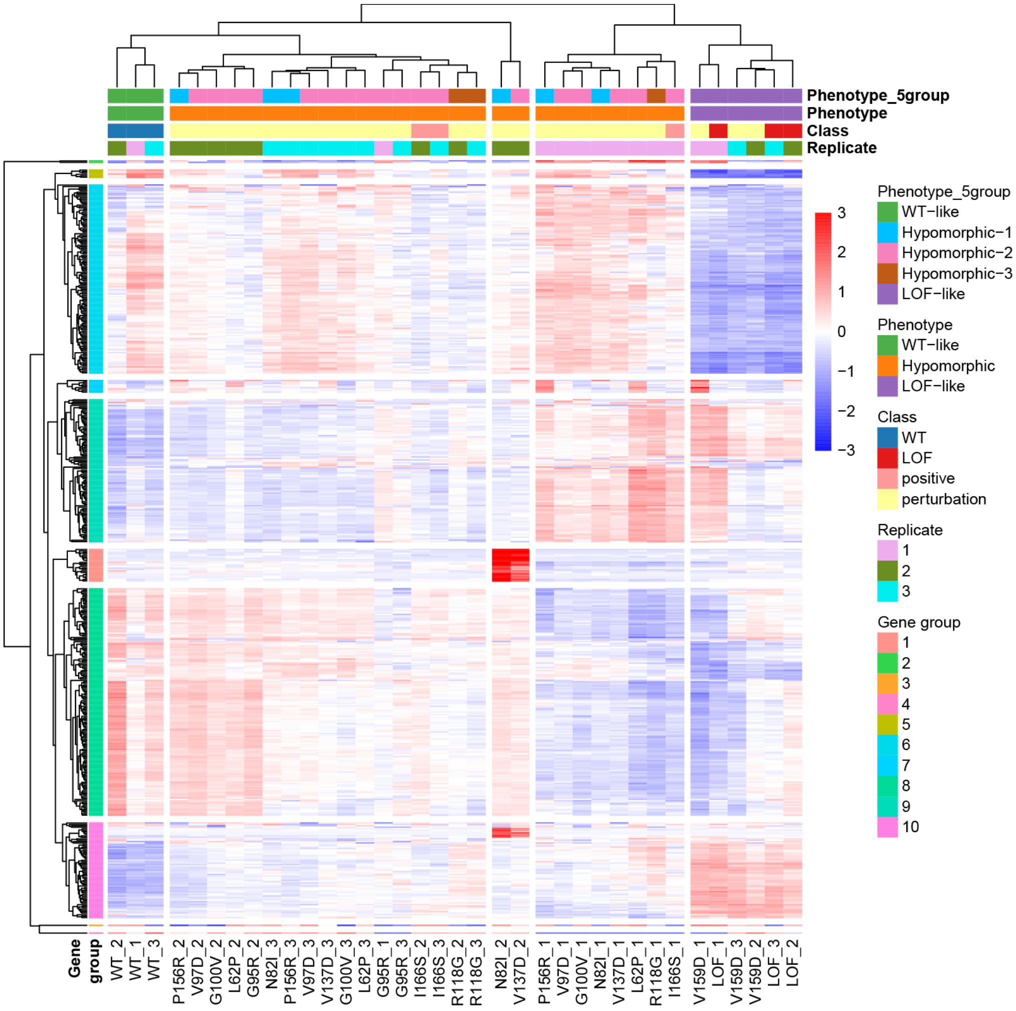


**Supplementary Figure 11. Hierarchical clustering of bulk RNA-seq samples before batch effect removal between replicates.**

Hierarchical clustering of samples (rows) and genes (columns), using top 2000 variable genes obtained from scRNA-seq. All sample replicates are present. Replicates 2 of samples with N82I and V137D mutations (N82I_2 and V137D_2), are identified as outliers and subsequently removed before batch effect removal between replicates and downstream analyses. The leaves of the variant dendrogram are ordered by increasing $T2_{WT}$ scores. Gene expression values are z-scored.

### SUPPLEMENTARY TABLES

**Supplementary Table 1.** Library of 117 variants, corresponding variant class, amino acid substitution, codon change, VEST and FoldX scores, and predicted binding partners for each residue.

**Supplementary Table 2.** Library of 112 variants that are selected for downstream analysis after filtering, their phenotypic annotations, $T2_{WT}$ and $T2_{LOF}$ scores, respective p-values, fitness scores, VEST and FoldX scores, number of occurrences in COSMIC and MLL datasets, binding information for DNA and CBFB, selected for validation or not.

**Supplementary Table 3.** Cluster enrichment of single cells (unsupervised clusters from Figure 2a), for each perturbation variant, based on log of odds ratios obtained using Fisher's exact test. Positive values indicate enrichment, while negative values indicate depletion.

**Supplementary Table 4.** Gene group (Figure 3a) scores for each phenotype cluster.

**Supplementary Table 5.** Gene set overrepresentation analysis results for GO Biological Process terms for each gene program (Figure 3a).

**Supplementary Table 6.** Gene set overrepresentation analysis results for Reactome pathways for gene markers of each pattern (Supplementary Figure 7).

**Supplementary Table 7.** Classifier predictions for all possible RUNX1 Runt domain variants.

**Supplementary Table 8.** Gene set overrepresentation analysis results for Reactome pathways for each gene group (Figure 6).

**Supplementary Table 9.** Gene set overrepresentation analysis results for Reactome pathways for each gene group (Supplementary Figure 10).

**Supplementary Table 10.** Primers.
